# Multimodal profiling resolves the functional and spatial organization of colorectal cancer liver metastases

**DOI:** 10.64898/2026.11.18.767576

**Authors:** Maelle Picard, Arnaud Guille, Emilie Denicolai, Nadiya Belfil, Pascal Finetti, Claire Germier, Gwenaël Lumet, Bernadette de Rauglaudre, Etienne Dougy, François Bertucci, David Jeremie Birnbaum, Emilie Mamessier

## Abstract

**Background:** Colorectal cancer (CRC) is the second leading cause of cancer-related mortality, largely because of liver metastases (LM). Despite advances in perioperative treatment and surgery, recurrence after resection of CRC liver metastases (CRC-LM) remains common. The cellular complexity of treatment-exposed CRC-LM and the interactions among malignant, stromal, immune, vascular and liver-resident populations remain incompletely understood, limiting the identification of biological vulnerabilities associated with disease progression and recurrence.

**Methods:** We used single-nuclei RNA sequencing (snRNAseq) to characterize the cellular composition and transcriptional states of nine chemotherapy-treated CRC-LM specimens. Single-nucleus profiles were integrated with spatial transcriptomic and protein-based analyses to define the functional and spatial organization of the metastatic ecosystem. We further compared tumors from patients with early versus no or later relapse and investigated candidate cell-cell interactions associated with recurrence-related cellular states.

**Results:** Single-nucleus profiling recovered malignant epithelial tumor cells, fibroblasts, immune cells, endothelial cells and liver resident epithelial populations across samples and revealed substantial functional heterogeneity within each compartment. Malignant epithelial cells occupied interconnected stem-like, proliferative, differentiated and invasive states, whereas stromal, immune and vascular populations exhibited distinct transcriptional programs organized within specific tissue niches. Comparative analysis identified a fibroblast subtype enriched in early-relapse metastases, which was reproducibly identified in independent LM-CRC single-cell datasets. This population showed features consistent with a role in tumor dissemination and appeared to actively shape the metastatic microenvironment.

**Conclusion:** Our study provides an integrated single-nucleus and spatial characterization of the cellular organization of chemotherapy-treated CRC-LM. It reveals functionally specialized multicellular niches and identifies a fibroblast state associated with early relapse and epithelial and endothelial plasticity.

## Introduction

Metastatic disease accounts for most cancer-related deaths and remains the principal cause of mortality in CRC patients worldwide [1,2]. The liver is the most frequent site of CRC metastasis [3,4], affecting more than 60% of patients with metastatic disease and the majority (83%) of patients who ul-timately die from CRC [5]. Surgical resection remains the only curative option, but it is feasible in only approximately 20% of patients, [6–8], and recurrence within two years is common even after apparent complete removal of the metastatic lesions [2,9]. This clinical scenario underscores the need to gain a deeper understanding of the biological mechanisms and cell-cell interactions supporting the early estab-lishment, progression and recurrence of CRC-LM.

CRC-LM are complex ecosystems formed by genetically and phenotypically diverse malignant cell populations that crosstalk with other cellular compartments such as immune cells, fibroblasts, vascular cells and liver resident cells. This diverse ecosystem is characterized by unique cell-cell interactions with-in spatially organized tissue niches that are further influenced and functionally reprogrammed by me-chanical tissue forces and exposure to environmental stressors such as inflammatory cues and systemic therapy [6,10]. Their composition and functional state may influence metastatic growth, treatment re-sponse, and subsequent relapse. Understanding CRC-LM therefore requires approaches that capture not only the diversity of their cellular components, but also their spatial organization and functional inter-play.

Single-cell transcriptomic studies have begun to reveal the cellular heterogeneity of CRC-LM, including extensive remodeling of the immune, malignant and stromal compartments [11]. More recent studies integrating single-cell and spatial transcriptomics, epigenomic profiling, metabolomics or multiplex pro-tein imaging have further delineated the spatial and functional organization of specific cellular states in the metastatic liver [12]. However, prior work has focused on specific cellular compartments, and a comprehensive portrayal of how metastatic tumor cells are jointly organized amidst hepatic, immune, stromal and vascular populations is still lacking. Moreover, the relationship between the composition of treatment-exposed metastatic ecosystems and subsequent clinical outcomes, including the timing of relapse, remains poorly defined. A more integrated characterization of these environments could uncov-er biological vulnerabilities, and candidate features relevant to patient stratification and the clinical management of metastatic CRC.

Here, we combined single-nucleus RNA sequencing (snRNAseq), spatial transcriptomics and proteomic analyses to characterize the cellular and functional heterogeneity of resected CRC-LM. snRNAseq is par-ticularly suited to fibrotic, necrotic, and treatment-exposed tissues, as it reduces biases introduced by tissue dissociation and facilitates the recovery of malignant, hepatic, stromal, vascular, and immune compartments [13]. We integrated these transcriptional profiles with spatial information to determine how distinct cellular states are organized within metastatic lesions and with proteomic data to validate the biological programs identified. Using this multimodal framework, we generated an integrated map of CRC-LM and explored differences between tumors from patients experiencing early relapse or not, iden-tifying a fibroblast state associated with early metastatic relapse.

## Results

### Single-nuclei profiling captures the broad cellular composition of colorectal cancer liver metastases

To define the cellular composition of CRC-LM, we performed snRNAseq on surgically resected specimens (n=9) from patients who had undergone chemotherapy treatment (Figure 1A; see Methods). Following quality control (*see Methods*), we obtained transcriptional profiles for a total of 31,507 high-quality nuclei, hereafter referred to cells, which clustered into six major malignant and non-malignant cellular compartments (Figure 1B). Clusters were annotated based on the expression of canonical markers genes (Figure 1C). All major compartments were represented across all patients, although their relative abundance varied between individual metastases (Figure 1D). This interpatient variation underscored the cellular complexity of the metastatic ecosystem and provided the basis for a more detailed analysis of the transcriptional states contained within each compartment. Cell count (Figure 1E) and proportions (Figure 1F) revealed that resident liver cells was the most abundant (9,837 cells for hepatocytes, 1,952 cells for cholangiocytes, for a total of 37.4% of cells), followed by epithelial tumor cells (6,420 cells, 20.4%), fibroblasts (5,807 cells, 18.4%), immune cells (4,825 cells, 15.3%), and endothelial cells (2,666 cells, 8.5%).

**Figure 1:**
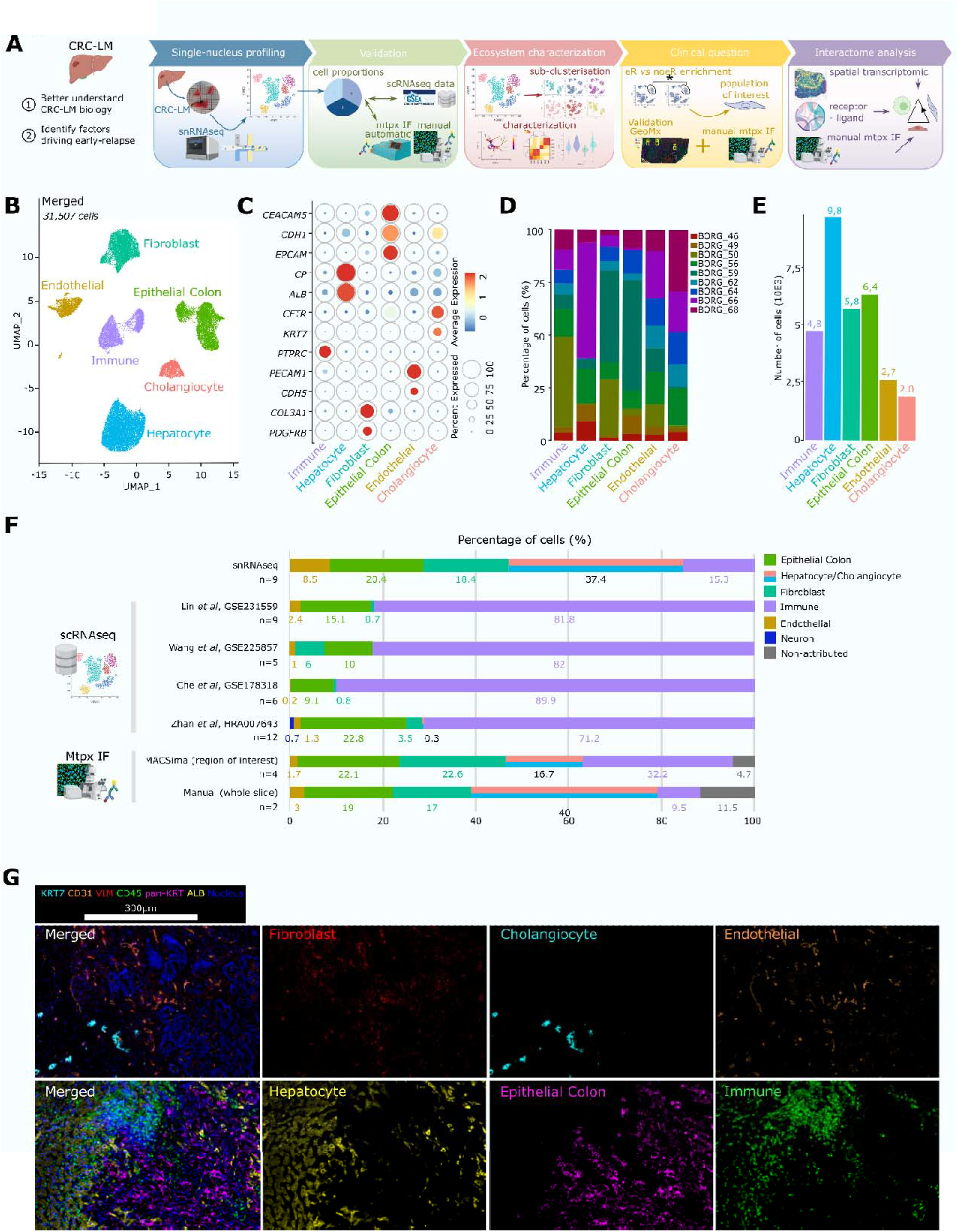
Characterization and validation of the major cellular compartments of CRC-LM profiled by snRNAseq. (A) General workflow of the study. (B) Uniform Manifold Approximation and Projection (UMAP) of the major cellular compartments obtained from 9 samples of CRC-LM analyzed by snRNAseq. (C) Dotplot representing the expression of canonical markers genes used to annotate major cellular compartments. Dot size is for percent of cells expressing the gene while color scale is for relative expression intensity. (D) Barplot representing the patient’s contribution for each major cellular compartment. (E) Barplot representing the absolute count for each major cellular compartment. (F) Barplot representing the proportions of major cellular compartments obtained by different approaches: snRNAseq referred to our data while Lin, Wang, Che, and Zhan lines referred to scRNAseq publicly available datasets collected (*see Methods*). Accession number were mentioned for scRNAseq datasets. Two last lines referred to cellular proportions obtained by mtpx IF performed either automatically using the MACSima technology on region of interest and manually using whole slide spectral immunofluorescence (*see Methods*). n: effective. (G) Microscopy images of the major cellular compartments detected at the protein level by manual multiplexed immunofluorescence.

Because tissue dissociation can introduce biases in the recovery of specific cell types, we next compared the cellular representation obtained by our snRNAseq approach with that reported in published single-cell RNA sequencing datasets (scRNAseq) of CRC-LM (*see Methods*). Dissociation-based scRNAseq displayed global depletion of resident liver cells (<1%), and poor representation of fibroblasts and endothelial cells (6% and 2.4% of total cells maximum, respectively), with epithelial tumor cells accounting for 9% to 23% of cells, while immune cells constituted the dominant compartment, representing more than 70% of the recovered cells. By contrast, single-nucleus profiling recovered a broader representation of malignant epithelial, mesenchymal, vascular, immune, and liver-resident populations (Figure 1F). Consistently, multiplexed immunofluorescence (mtpx IF) of our samples revealed that fibroblasts represented 22.6% of cells, epithelial tumor cells 22.1%, healthy liver cells 16.7%, endothelial cells 1.7%, immune cells 32.2%, while 4.7 % of cells could not be assigned to a specific compartment (Figure 1F, Supplementary Figure 1B). Given the potential bias introduced by the selection of the region of interest (ROI), whole slide manual mtpx IF was performed to determine cell proportions on two additional patients. Fibroblasts accounted for 17% of cells, endothelial cells for 3%, epithelial tumor cells for 19%, immune cells for 9.5%, liver-resident cells for 40% (Figure 1F-G). Altogether, these data show that snRNAseq reduces some of the recovery biases associated with dissociation of fibrotic and treatment-exposed metastatic tissues and provides a faithful representation of the LM environment.

### Malignant epithelial cells span a continuum of differentiation and proliferation states

Despite prior exposure to chemotherapy, we isolated 6,420 high-quality epithelial tumor nuclei, which reclustered into seven transcriptionally distinctive clusters (from EpiColon_c0 to EpiColon_c6, Figure 2A). Most clusters were shared across patients, except for EpiColon_c6 which was detected on a single patient and was thus removed from subsequent analyses (Figure 2B). The malignant identity of epithelial clusters was inferred from copy number variation (CNV) relative to hepatocytes (Figure 2C).

**Figure 2:**
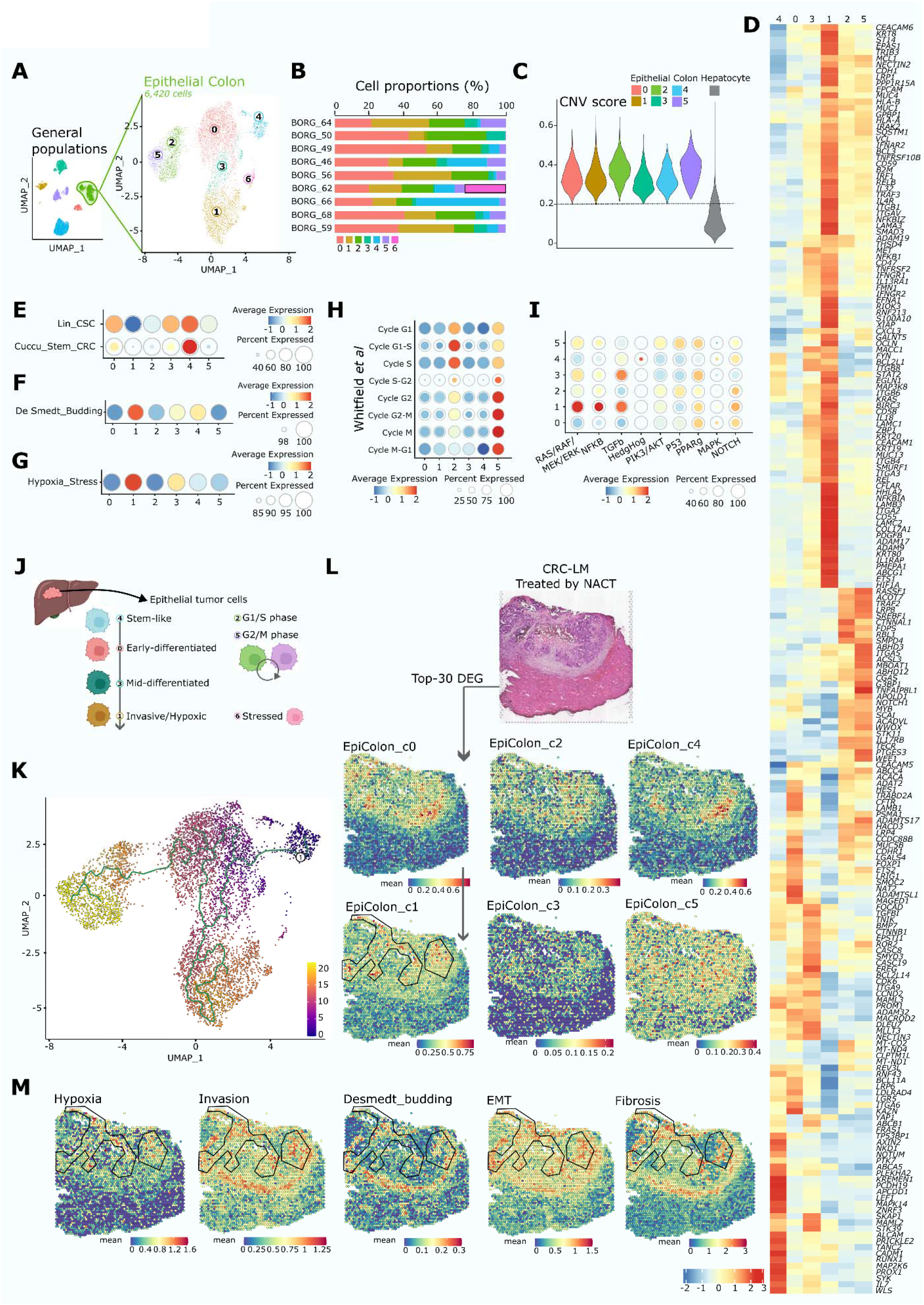
Characterization of the epithelial tumoral compartment of CRC-LM. (A) UMAP of epithelial tumor clusters obtained by reclustering the epithelial main cellular compartment. (B) Barplot representing the patient’s contribution for each epithelial tumor cluster. A frame highlights the cluster EpiColon_c6 coming from only one patient. (C) Violin plot representing the copy number variation of epithelial tumor clusters compared to the hepatocytes compartment as a reference. (D) Heatmap representing the expression of DEG of interest across epithelial tumor clusters and used for their annotation. Color scale is for relative expression intensity. (E) Dot plot representing the enrichment of CSC signatures across epithelial tumor clusters. Dot size is for percent of cells expressing the signatures while color scale is for relative expression intensity. (F) Dot plot representing the enrichment of budding signature across epithelial tumor clusters. Dot size is for percent of cells expressing the signature while color scale is for relative expression intensity. (G) Dot plot representing the enrichment of hypoxia/stress signature across epithelial tumor clusters. Dot size is for percent of cells expressing the signature while color scale is for relative expression intensity. (H) Dot plot representing the enrichment of cell cycle signatures across epithelial tumor clusters. Dot size is for percent of cells expressing the signatures while color scale is for relative expression intensity. (I) Dot plot representing the enrichment of main signaling pathway expressed by CRC cells across epithelial tumor clusters. Dot size is for percent of cells expressing the signatures while color scale is for relative expression intensity. (J) Graphical summary of the CRC-LM epithelial tumor cell composition and annotations. (K) Pseudotime representation of the trajectory inference from EpiColon_c4 stem-like cluster (defined as origin) toward other epithelial tumoral clusters using Monocle3 method (see Methods). (L) Projection of the expression of epithelial tumor clusters metagenes (top-30 DEG) on a spatial transcriptomic data from CRC-LM treated by neoadjuvant chemotherapy (NACT). Color scale is for absolute expression intensity. Arrows indicate the reading direction. Polygonal frame highlights the region enriched for EpiColon_c1. (M) Projection of the expression of hypoxia, invasion, budding, EMT and fibrosis signatures on a spatial transcriptomic data from CRC-LM treated by NACT. Color scale is for absolute expression intensity. Polygonal frame highlights the region enriched for EpiColon_c1.

To define the biological states represented by these malignant epithelial populations, we integrated cluster-specific differentially expressed genes (DEG) (Figure 2D, Supplementary Table 1-2) with gene ontology, pathway, and established transcriptional signatures (Figure 2E-I). Signatures tested are listed in Supplementary Table 3. These analyses revealed four major features of epithelial heterogeneity comprising stem-like, differentiating states, proliferative states, and an invasive/hypoxic phenotype.

EpiColon_c4 displayed the strongest enrichment in stemness, including genes involved in Wnt signaling and high scores in previously described cancer stem cell (CSC) signatures (Figure 2E). Stemness features were detected less prominently in EpiColon_c0 and EpiColon_c3, suggesting a progressive loss of stem-like features in these populations. EpiColon_c0 was enriched in genes implicated in mucus secretion, such as MUC5B which is associated with immature secretory cells in CRC [14]. EpiColon_c3 exhibited features compatible with a more differentiated state, including apoptosis and BMP-associated programs, and genes involved in TGFβ signaling (Figure 2D, Supplementary Table 1-2). Based on these transcriptional states, EpiColon_c4, EpiColon_c0, and EpiColon_c3 were annotated as stem-like, early-differentiated, and mid-differentiated epithelial states, respectively (Figure 2J).

By contrast, EpiColon_c1 was characterized by a distinctive phenotype retaining canonical epithelial features and was also enriched for markers of cells associated with the invasive front, extracellular matrix (ECM) remodeling, hypoxia and TGFβ signaling. RAS/RAF/MEK/ERK and NF-kβ signaling, were also preferentially represented in this cluster, along with enrichment in a signature representative of budding and hypoxic/stressed phenotypes (Figure 2F-G, I, Supplementary Table 1-2), altogether supporting its annotation as an invasive/hypoxic phenotype. Anti-apoptotic programs were also represented in this cluster (along with EpiColon_c5), may further contribute to the persistence of these cells under adverse microenvironmental conditions (Figure 2D, Supplementary Table 1-2).

In parallel, EpiColon_c2 and EpiColon_c5 formed a distinct proliferative branch. Cell-cycle signatures indicated that EpiColon_c2 was predominantly associated with G1/S progression, while EpiColon_c5 displayed a G2/M-phase program (Figure 2H, Supplementary Table 1-2). DEG and pathways analysis revealed that both populations were enriched for DNA replication and repair, chromatin remodeling, topoisomerase activity, mitotic-spindle organization, kinetochore function, and chromosome segregation. Their transcriptional differences were consistent with their respective positions within the cell cycle, with EpiColon_c2 showing greater representation of epigenetic-regulatory programs, and EpiColon_c5 being enriched for, G2/M progression and lipid-metabolic pathways, mirroring the biosynthetic requirements of proliferating cells (Supplementary Figure 3B-C). Consistently, PI3K-Akt and p53-associated programs, were highly represented in these cycling states (Figure 2I).

Pathway enrichment analysis across the epithelial tumor clusters revealed marked heterogeneity in signaling programs (Figure 2I). Hedgehog signaling was enriched in stem_like_EpiColon_c4, whereas PPARγ signaling, which is associated with cellular differentiation, was detected across most epithelial states but was comparatively limited in EpiColon_c4, consistent with its stemness phenotype. NOTCH activity, which is implicated in differentiation, was broadly represented across all malignant epithelial populations, whereas TGFβ was most prominently detected in EpiColon_c1 and EpiColon_c3, consistent with epithelial plasticity along the transition toward the invasive/hypoxic state. Together, these patterns indicate that the epithelial clusters represent overlapping biological states exhibiting features compatible with differentiation, proliferation, stress adaptation, and microenvironmental signaling rather than completely discrete transcriptional identities.

In addition, discrete epithelial clusters expressed candidate neoantigens (EpiColon_c0), therapeutic targets (EpiColon_c0, 1, 3 and 5) and oncogenes/tumor suppressors genes (EpiColon_c0, 2, 5) (Supplementary Table 2). For example, EpiColon_c1 expressed *CD47* which binds to SIRPα in macrophages leading to immune escape [15]. Other populations expressed mediators with potential paracrine functions, including the neutrophil chemoattractant [16] *CXCL3* in EpiColon_c3, *IL17RB* in EpiColon_c2, and the prostaglandin-synthesis enzyme PTGES3 in EpiColon_c5. Altogether, these data support the notion that the transcriptionally distinct and diverse malignant states along a differentiation continuum contribute to remodeling of the metastatic microenvironment (Figure 2J).

The relationships among these epithelial states suggested that they may represent interconnected positions along defined phenotypic trajectories. We therefore performed trajectory inference taking the stem-like EpiColon_c4 population as the putative origin. Based on this analysis, EpiColon_c0 was the earliest differentiated subtype and subsequently diverging toward a proliferative lineage (EpiColon_c2 then EpiColon_c5 clusters) or a progressive differentiation (EpiColon_c3 then EpiColon_c1 clusters). Slingshot trajectory revealed that a subset of EpiColon_c1 cells was also positioned along trajectories approaching proliferative states, suggesting potential plasticity between invasive and cycling phenotypes (Figure 2K, Supplementary Figure 3D). Therefore, and consistent with previous analyses, rather than defining a linear hierarchy, the trajectory analysis supports a continuum of plastic malignant states encompassing stemness, differentiation, proliferation, and invasion.

We next asked whether these transcriptional states were associated with specific spatial niches by mapping their metagene signatures in our CRC-LM samples. Stem-like EpiColon_c4 and early-differentiated EpiColon_c0 showed relatively restricted distributions, consistent with their localization within discrete tumor niches. In contrast, EpiColon_c2 and EpiColon_c3 were more broadly distributed within the tumor core. The proliferative EpiColon_c5 signature was detected in the tumor, but also in the non-malignant tissue, highlighting a classical proliferative function (Figure 2L). Notably, the invasive/hypoxic EpiColon_c1 signature localized preferentially to hypoxic tumor regions overlapping with transcriptional programs associated with invasion, budding, epithelio-mesenchymal transition (EMT) and fibrosis (Figure 2M, Supplementary Table 4). This spatial context was consistent with its transcriptional phenotype and positioned EpiColon_c1 within tissue regions potentially permissive to tumor cell migration and dissemination.

Collectively, these analyses expose substantial epithelial heterogeneity within CRC-LM epithelial cells and support the co-existence of interconnected malignant states spanning stemness, proliferation and invasive capacity that may be shaped and actively modulate the spatial organization of the metastatic tissue environment.

### Mesenchymal populations are characterized by defined functional and spatial tissue niches

Fibroblasts are key stromal cell types and play a central role in tumor initiation and progression [17,18]. However, the identity and diversity of this heterogeneous and highly plastic population keep being refined in CRC-LM. Reclustering of the mesenchymal compartment resolved eight populations comprising transcriptionally distinct fibroblast states, pericytes and vascular smooth muscle cells (VSMC) (from Fibro_c0 to Fibro_c7), which were shared by most patients in variable proportions (Figure 3A-B). Annotations were established regarding canonical markers and pathways identified in DEG (Supplementary Table 5-6) and enrichment in previously described fibroblast signatures and markers (Figure 3D-J). It allowed us to relate these populations to established immunoregulatory and myofibroblast states and additional transcriptional heterogeneity within this cellular compartment. Pericytes and VSMC were identified in Fibro_c5 and Fibro_c6 respectively by their expression in canonical markers (Figure 3C).

**Figure 3:**
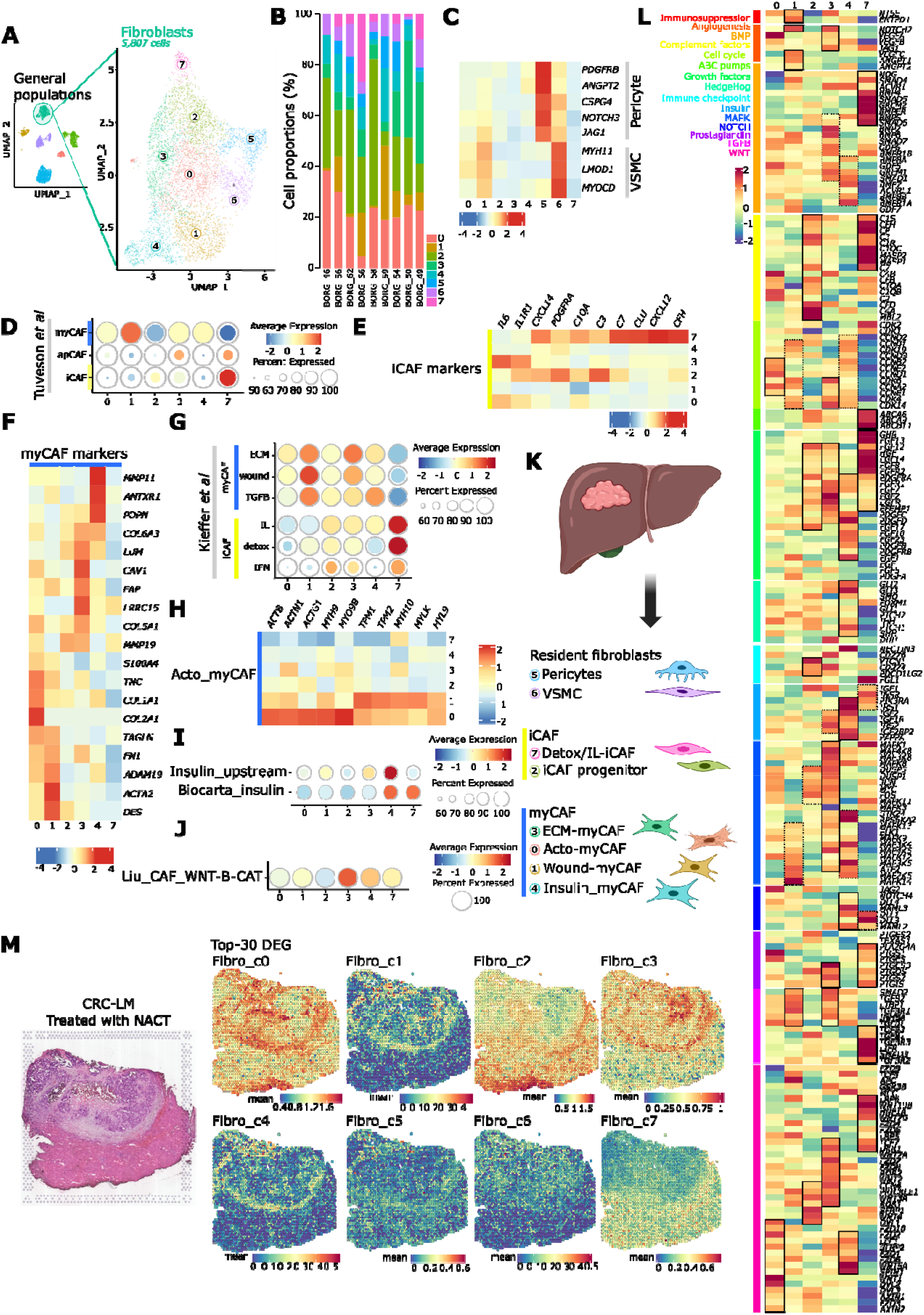
Characterization of the fibroblast compartment of CRC-LM (A) UMAP of the fibroblast clusters obtained by reclustering of the fibroblast major cellular compartment. (B) Barplot representing the patient’s contributions for each fibroblasts cluster. (C) Heatmap representing the expression of canonical markers genes for pericytes and VSMC. Color scale if for relative expression intensity. (D) Dotplot representing the expression of broad CAF signatures across fibroblasts clusters. Dot size is for percent of cells expressing the signatures while color scale is for relative expression intensity. A blue frame was attributed to myCAF while yellow frame to iCAF to facilitate further readability. (E) Heatmap representing the expression of main iCAF markers across fibroblasts clusters. Color scale if for relative expression intensity. (F) Heatmap representing the expression of main myCAF markers across fibroblasts clusters. Colo scale if for relative expression intensity. (G) Dot plot representing the expression of myCAF and iCAF subtypes signatures across fibroblasts clusters. Dot size is for percent of cells expressing the signatures while color scale is for relative expression intensity. (H) Heatmap representing the expression of the main acto-myCAF markers genes across fibroblasts clusters. Color scale is for relative expression intensity. (I) Dot plot representing the expression of insulin pathway signatures across fibroblasts clusters. Dot size is for percent of cells expressing the signatures while color scale is for relative expression intensity. (J) Dot plot representing the expression of CAF_Wnt/β-Catenin signature across fibroblasts clusters. Dot size is for percent of cells expressing the signature while color scale is for relative expression intensity. (K) Graphical summary of the CRC-LM fibroblasts composition and annotations. (L) Heatmap representing the expression of DEG across fibroblasts clusters, grouped by gene ontologies and used for their annotation. Color scale is for relative expression intensity. Solid line frames highlight the highest expression while dash line frames are for mid expression. (M) Projection of the expression of fibroblasts clusters metagenes (top-30 DEG) on a spatial transcriptomic data from CRC-LM treated by NACT. Color scale is for absolute expression intensity.

The presence of cancer-associated fibroblasts (CAF) consensual subtypes, namely immunoregulatory CAF (iCAF), myofibroblastic CAF (myCAF), and antigen-presenting CAF (apCAF), were evaluated using gene signature, excluding pericytes and VSMC, which were already annotated. Tuveson et al’s i/my/apCAF broad identification revealed that myCAF signature was strongly enriched in Fibro_c1 cluster and, to a lesser extent, in Fibro_c0, Fibro_c3 and Fibro_c4 clusters. Approximately 80% of the cells from Fibro_c3 and Fibro_c7 clusters harbored an apCAF signature enrichment, and Fibro_c7 cluster was clearly enriched in iCAF signature. Also, 80% of the cells from Fibro_c2 cluster harbored an iCAF signature enrichment suggesting that both Fibro_c7 and Fibro_c2 correspond to iCAF (Figure 3D). To confirm, we examined the expression of established iCAF markers reported in the literature. Fibro_c2 and Fibro_c7, highlight an increased expression of genes related to inflammation and complement cascade, further validating the iCAF annotation (Figure 3E). Similarly, Fibro_c0, c1, c3 and c4 clusters were enriched in several myCAF markers (Figure 3F), despite substantial heterogeneity in marker expression. To further refine this analysis, we applied the Kieffer et al’s well-established i/myCAF subclassification (Figure 3G). ECM_myCAF and wound_ECM_myCAF were strongly expressed in both Fibro_c1 and Fibro_c3. TGFβ-myCAF signature was enriched in both Fibro_c1 and Fibro_c4 at similar average expression levels, suggesting a variable enrichment likely due to redundancy in myCAF features (Supplementary Figure 3A). Regarding iCAF subtypes, Fibro_c7 displayed strong enrichment for IL-iCAF and detox-iCAF signatures, with additional moderate enrichment for IFNγ-iCAF, shared with Fibro_c2 (Figure 3G). It was thus annotated as a mix of the two first iCAF subtypes. Fibro_c2 was characterized by only 38 DEG, including genes coding for complement cascade, HLA or IL1R1 explaining the IFNγ-iCAF enrichment (Supplementary Table 5-6). However, its low number of DEG compared to the other fibroblasts subtypes could suggest a basal phenotype as an iCAF progenitor profile. By contrast, Fibro_c0 did not reveal any clear enrichment in these CAF signatures (Figure 3G). As the signature of acto-myCAF - another CAF subtype well defined in the literature as involved in cell contractility - was not available, actin and myosin-related gene expression was examined. Most genes were predominantly enriched in Fibro_c0 - and in Fibro_c1 consistently to their involvement in wound healing responses (Figure 3H). Considering levels of expression of specific markers, we found that Fibro_c4 exhibits a specific and marked enrichment in insulin receptor signaling which was confirmed by signature enrichment (upstream: *IGF1R, IRS1/2, PAPPA, IGF2BP2, PIK3R1, IGF2*, Biocarta: complete signaling pathway) (Figure 3I, L, Supplementary Table 6). Finally, we found a strong enrichment in Fibro_c3 of CAF_Wnt/β-Catenin signature (Figure 3J).

Altogether, these analyses led to the following annotations: Fibro_c0 as acto-myCAF, Fibro_c1 as wound-myCAF, Fibro_c3 as ECM-myCAF, Fibro_c5 as pericytes, Fibro_c6 as VSMC, and Fibro_c7 as detox-/IL-iCAF while Fibro_c2 was defined as an iCAF progenitor and Fibro_c4 did not align with the i/myCAF framework suggesting a new insulin-related myCAF subtype (Figure 3K).

To define the functional specialization of fibroblast states in CRC-LM, we performed gene ontology and pathway enrichment analyses using cluster-specific DEG (Figure 3L), after excluding broadly shared fibroblast programs, notably those related to cell adhesion and ECM synthesis and remodeling. Wnt signaling pathway was shared across fibroblasts clusters but was more prominently expressed in ECM-myCAF/Fibro_c3, followed by acto-myCAF/Fibro_c0, iCAF-progenitor/Fibro_c2, insulin_myCAF/Fibro_c4, and detox-IL-iCAF/Fibro_c7. Similarly, TGFβ signaling characterized several fibroblast states, but was particularly enriched in wound-myCAF/Fibro_c1, ECM-myCAF/Fibro_c3 and detox-IL-iCAF/Fibro_c7. Growth factor related programs were distributed across iCAF-progenitor/Fibro_c2, ECM-myCAF/Fibro_c3, insulin_myCAF/Fibro_c4, detox-IL-iCAF/Fibro_c7; whereas MAPK signaling was enriched in ECM-myCAF/Fibro_c3 mainly, then by wound-myCAF/Fibro_c1, iCAF-progenitor/Fibro_c2 and insulin-myCAF/Fibro_c4. These shared programs superimposed on more cluster-restricted features may reflect a functional specialization of diverse fibroblast populations within the metastatic environment.

Wound-myCAF/Fibro_c1 displayed prominent expression of angiogenesis and tumor-stroma communication programs and genes implicated in adenosine-mediated immunosuppression, a mechanism previously linked to metastatic progression [19]. Additionally, this cluster were characterized by DEG including TAGLN, POSTN and FAP, all of which are TGFβ-related factors previously associated with CRC-LM outgrowth [20] (Supplementary Table 5-6).

ECM-myCAF/Fibro_c3 showed the most striking enrichment of signaling and remodeling programs. In addition to angiogenic, growth factors, prostaglandin, TGFβ, MAPK, and Wnt-related, this fibroblast population expressed genes linked to IL1 signaling (IL1R1) and hypoxic responses (HIF1A). This cluster also expressed FAP and INHBA, CAF-associated markers that have been linked to adverse outcome in gastric cancer [21], as well as ITGBL1, which has been reported to promote formation of fibroblast-rich metastatic niches through extracellular-vesicle-mediated signaling [22] (Supplementary Table 5-6). Together, these data indicate that ECM-myCAF/Fibro_c3 represents a highly activated fibroblast state combining ECM remodeling with microenvironment-oriented signaling.

Insulin-myCAF/Fibro_c4 was distinguished by enrichment of insulin, Hedgehog, MAPK, NOTCH, growth-factor, and WNT signaling programs. This population expressed NECTIN3, encoding for a ligand of the inhibitory receptor TIGIT, as well as PDPN, a CAF marker previously associated with poor prognosis [23]. This cluster further expressed PDGFRA and ITGA11, which previous work has reported in a fibroblast population present in early-stage bladder cancer and proposed to interact [24]. Lastly, we noted high ANTXR1 levels in insulin-myCAF/Fibro_c4, which has been associated with immunotherapy resistance, early relapse, and poor prognosis in ovarian cancer [25] (Supplementary Table 5-6). Together these findings point to a myCAF subtype enriched in insulin pathway.

The iCAF-progenitor/Fibro_c2 population displayed PDGFRA expression, consistent with an inflammatory fibroblast identity [26], and was enriched for complement-related genes together with detox-IL-iCAF/Fibro_c7. iCAF-progenitor/Fibro_c2 cells also showed preferential expression of immune-checkpoint-related genes (Supplementary Table 5-6), suggesting a potential role on immune regulation and tumor-stroma crosstalk.

Detox-IL-iCAF/Fibro_c7 exhibited a distinct inflammatory and stress-adaptation profile. Consistent with its annotation, this cluster showed enrichment of ATP-binding cassette transporters and was characterized by BMP-, complement cascade-, prostaglandin-, TGFβ-related programs. Detox-IL-iCAF/Fibro_c7 also expressed CXCL12, consistent with a potential role on immune-cell recruitment and T-cell regulation [27]. Notably, it expressed FGL1, a major ligand of the immune checkpoint LAG3, involved in T cell inhibition [22,28], as well as HGF that has been shown to enhance tumor growth in intrahepatic cholangiocarcinoma models [29] (Supplementary Table 5-6).

Finally, acto-myCAF/Fibro_c0 was characterized predominantly by contractile and proliferative features.

The functional specialization inferred from these transcriptional programs was mirrored by the spatial organization of the fibroblast and mural populations. We applied spatial transcriptomics to map cluster-specific metagene signatures across metastatic tissues and found that acto-myCAF/Fibro_c0, wound-myCAF/Fibro_c1, ECM-myCAF/Fibro_c3 and insulin-myCAF/Fibro_c4 signatures were preferentially enriched in fibrotic regions surrounding malignant epithelial aggregates in the tumor margin and fibrotic areas within the tumor core. This distribution was consistent with the ECM-remodeling, angiogenic, and growth factor programs that characterized these fibroblast states. By contrast, Fibro_c5/pericytes and Fibro_c6/VSMC were more difficult to locate, given the resolution and the scattered nature of these populations. Of note, iCAF-progenitor/Fibro_c2 and detox-IL-iCAF/Fibro_c7 were predominantly confined to non-malignant liver tissue regions, suggesting a more resident fibroblast phenotype (Figure 3M). Collectively, these analyses indicate that fibroblast heterogeneity in CRC-LM extends beyond the canonical iCAF-myCAF dichotomy and support a more granular functional specialization within compartmentalized metastatic tissue niches in the liver.

### Immune infiltrates of CRC-LM are characterized by immunosuppressive and exhaustive states

To characterize the immune landscape of CRC-LM, we reclustered 4,825 immune nuclei into ten populations (Immune_c0 to c9), which were represented at variable frequencies across patients (Figure 4A-B). A first DEG overview revealed two major lineages: a myeloid compartment (*HLA, ITGAM, ITGAX, CD163, CIITA etc*.), and a lymphoid compartment (*Immunoglobulin, PAX5, LCK, THEMIS, CD3 etc*.). Integration of well-established canonical markers, DEG, gene ontology analysis, and published immune-cell signatures subsequently resolved distinct resident, tumor-associated, and functionally altered immune states (Figure 4C-E and Supplementary Table 7-8).

**Figure 4:**
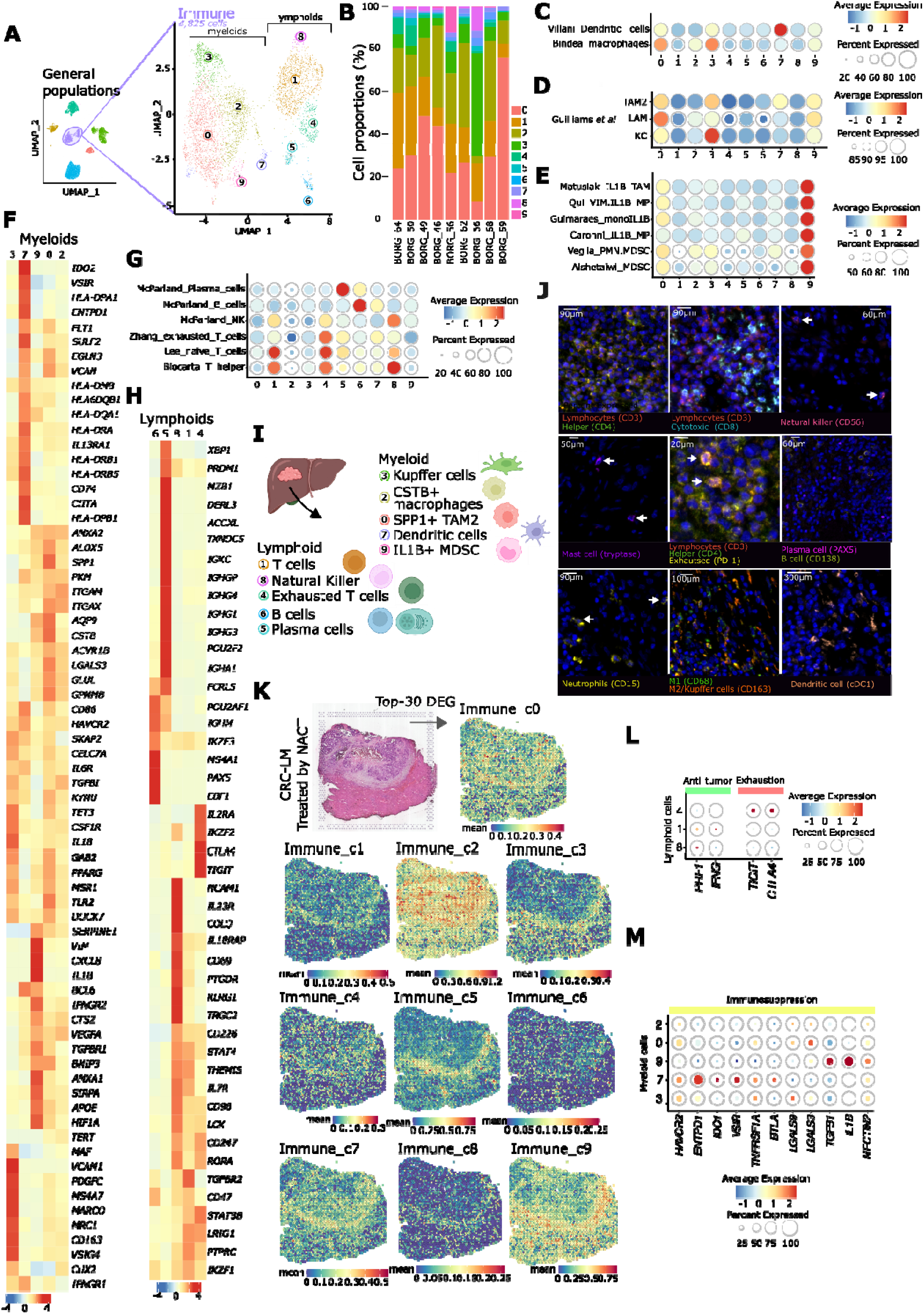
Characterization of the immune compartment of CRC-LM. (A) UMAP of immune clusters obtained by reclustering the immune main cellular compartment. Segregation between myeloid and lymphoid clusters was highlighted. (B) Barplot representing the patient’s contribution for each immune cluster. (C) Dot plot representing the enrichment of dendritic cell and macrophage signatures across immune clusters. Dot size is for percent of cells expressing the signatures while color scale is for relative expression intensity. (D) Dot plot representing the enrichment of TAM2, LAM and KC signatures across immune clusters. Dot size is for percent of cells expressing the signatures while color scale is for relative expression intensity. (E) Dot plot representing the enrichment of IL1β^+^ TAM and macrophages (MP), monocytes (mono) signatures and MDSC of polymorphonuclear (PMN) origin (or not) across immune clusters. Dot size is for percent of cells expressing the signatures while color scale is for relative expression intensity. (F) Heatmap representing the expression of DEG across myeloid immune clusters used used for their annotation. Color scale is for relative expression intensity. (G) Dot plot representing the enrichment of plasma cells, B cells, NK, and T cells signatures across immune clusters. Dot size is for percent of cells expressing the signatures while color scale is for relative expression intensity. (H) Heatmap representing the expression of DEG across lymphoid immune clusters used for their annotation. Color scale is for relative expression intensity. (I) Graphical summary of the CRC-LM immune cell composition and annotations. (J) Microscopy images validating the presence of most of the immune clusters at the protein level obtained by automatic mtpx IF. Arrows highlight the cells of interest. (K) Projection of the expression of immune clusters metagenes (top-30 DEG) on a spatial transcriptomic data from CRC-LM treated by NACT. Color scale is for absolute expression intensity. (L) Dot plot representing the enrichment of main genes involved either in anti-tumoral activity or T cell exhaustion across non-humoral lymphoid clusters. Dot size is for percent of cells expressing the genes while color scale is for relative expression intensity. (M) Dot plot representing the enrichment of immunosuppression-associated genes across myeloid clusters. Dot size is for percent of cells expressing the genes while color scale is for relative expression intensity.

The myeloid compartment comprised five populations (Immune_c0, c2, c3, c7, and c9). Immune_c0 and Immune_c3 both displayed macrophage signatures but represented distinct tumor-associated and liver-resident states. Immune_c0 was enriched for type-2 tumor-associated macrophage (TAM2) and lipid-associated macrophage (LAM) signatures, whereas Immune_c3 preferentially expressed Kupffer cell (KC) markers and signatures (Figure 4C-D). Immune_c2 displayed a more restricted macrophage program characterized by ITGAX and CTSB, consistent with a distinct CTSB^+^ myeloid population. Among these macrophage states, Immune_c0 exhibited the strongest tumor-associated and immunoregulatory phenotype. It expressed established TAM2-associated genes [30,31], angiogenic, cytokine-signaling, hypoxia-response, and metastasis-related programs. Notably, this population expressed SIRPA, the receptor for CD47 - which was expressed by EpiColon_c1 cluster - suggesting a potential immune escape axis. Immune_c0 further expressed CTSB, a marker defining a macrophage subtype responsible for the repression of memory immune hub in CRC-LM [32], as well as *KYNU, PLXDC2*, and *GPNMB* which expression have been associated with immunosuppression, stromal interactions, epithelial plasticity, and adverse outcomes [33–35]. Of note, expression of SPP1 further connected this population to macrophages interacting with FAP^+^ fibroblasts in primary and metastatic CRC [36,37] (Supplementary Table 8). In addition, Immune_c0 expressed hypoxia response programs [38,39], including some described in hypoxic and peri-necrotic glioblastoma [40], suggesting adaptation to metabolically and spatially restricted tumor niches.

Another myeloid state was represented by Immune_c9. This population showed enrichment in IL1β^+^ macrophage/monocytes signatures, as well as myeloid-derived suppressive cells (MDSC) associated programs (Figure 4E). IL1β^+^ myeloid populations have been associated with regulatory T cell recruitment and formation of an immunosuppressive tumor microenvironment [41] and colocalizing at the invasive tumor margin in kidney cancer [42] supporting annotation of Immune_c9 as an IL1β^+^ suppressive myeloid state. By contrast, the Immune_c7 cluster expressed canonical markers and signatures of dendritic cells (DC) (Figure 4C) and genes involved in tryptophan metabolism, adenosine-mediated immunosuppression consistent with an immunosuppressive DC phenotype (Supplementary Table 8).

The lymphoid compartment comprised B cells, plasma cells, NK cells, and heterogeneous T cell populations (Immune_c1, c4, c5, c6 and c8) (Figure 4G-H and Supplementary Table 7-8). Immune_c5 corresponded plasma cells exhibiting the class switch toward expression of immunoglobulin G and A, whereas Immune_c6 expressed canonical B cell markers and Immunoglobulin M transcripts, consistent with a less differentiated state. Immune_c8 showed strong NK cell signatures and canonical NK markers, although expression of T cell-associated genes revealed features of cytotoxic T cell populations, including γδ T cells. Thus, Immune_c8 was annotated as NK cells, keeping in mind a potential mix with other cytotoxic T cell populations.

Immune_c1 and Immune_c4 represented T cell enriched populations. Immune_c1 expressed signatures associated with helper T cell lineages, including a naive T cells signature, with Immune_c4 additionally expressing exhausted T cells signature and markers enrichment [43], supporting its annotation as an exhausted T cell population (Figure 4G, Supplementary Table 8).

We next evaluated whether these transcriptionally defined immune states occupied distinct spatial niches within chemotherapy-treated CRC-LM. The SPP1^+^ TAM2/LAM-like Immune_c0 population was detected both within the tumor core and in adjacent non-malignant liver regions, whereas the CTSB^+^ Immune_c2 population was preferentially localized within the tumor core. By contrast, Immune_c1, Immune_c5, Immune_c7, and Immune_c9 and to a lesser extent Immune_c8, were strongly enriched at the tumor margins, suggesting an immune exclusion. The localization of the IL1β^+^ suppressive myeloid Immune_c9 population at this interface was consistent with observations in another tumor type [42,43]. Kupffer Immune_c3 cells were distributed across the tumor margin and adjacent liver parenchyma, consistent with their resident hepatic identity. Immune_c4 and Immune_c6 showed no clear spatial pattern, potentially reflecting their relatively low abundance (Figure 4K).

The accumulation of several immune populations at the tumor margin raised the possibility that immune-cell localization was accompanied by impaired effector function. We therefore examined the expression of cytotoxic effector genes, exhaustion-associated receptors, and immunosuppressive mediators across lymphoid and myeloid populations (Figure 4L-M). Markers of exhaustion (*CTLA4* and *TIGIT*) were preferentially expressed by Immune_c4, whereas expression of genes encoding for effector molecules PRF1 and IFNG was relatively limited in lymphoid populations, consistent with reduced cytotoxic activity. In parallel, dendritic cells Immune_c7 and IL1β^+^ suppressive myeloid Immune_c9, and to a lesser extent the SPP1^+^ TAM2/LAM-like Immune_c0 population, expressed several immunoregulatory genes. Dendritic cells expressed *HAVCR2, ENTPD1, IDO1, VSIR, TNFRSF1A*, and *BTLA*, implicating pathways related to T cell inhibition, adenosine signaling, and tryptophan catabolism. Immune_c9 expressed TGFB1, IL1B, and NECTIN2, further supporting an immunosuppressive phenotype. The major immune populations identified transcriptionally were further supported at the protein level by mtpx IF in CRC-LM (Figure 4J). Granulocytes were detected by imaging (Supplementary Figure 3E) but were poorly represented in the single-nucleus dataset, likely because of their low RNA content and a strong ability to release RNAse [44], and further emphasizing the value of protein-based validation.

Collectively, the immune landscape of CRC-LM is characterized by co-existing myeloid and lymphoid states, including immunosuppressive and exhausted immune subsets, and a spatially organized immune-excluded tumor immune microenvironment.

### The vascular compartment contains an endothelial population with mesenchymal features

To characterize the vasculature compartment, we re-clustered 2,666 endothelial nuclei into eight subtypes (from Endo_c0 to Endo_c7, Figure 5A), all of which were shared across all patients, albeit in variable proportions (Figure 5B). To refine subtype identity, we integrated publicly available endothelial cell signatures (Figure 5C) and analyzed DEG with respect to canonical markers, gene ontologies and functional features. These analyses resolved populations associated with liver sinusoidal identity, vascular specialization, angiogenesis, and endothelial plasticity (Figure 5D and Supplementary Table 9-10).

**Figure 5:**
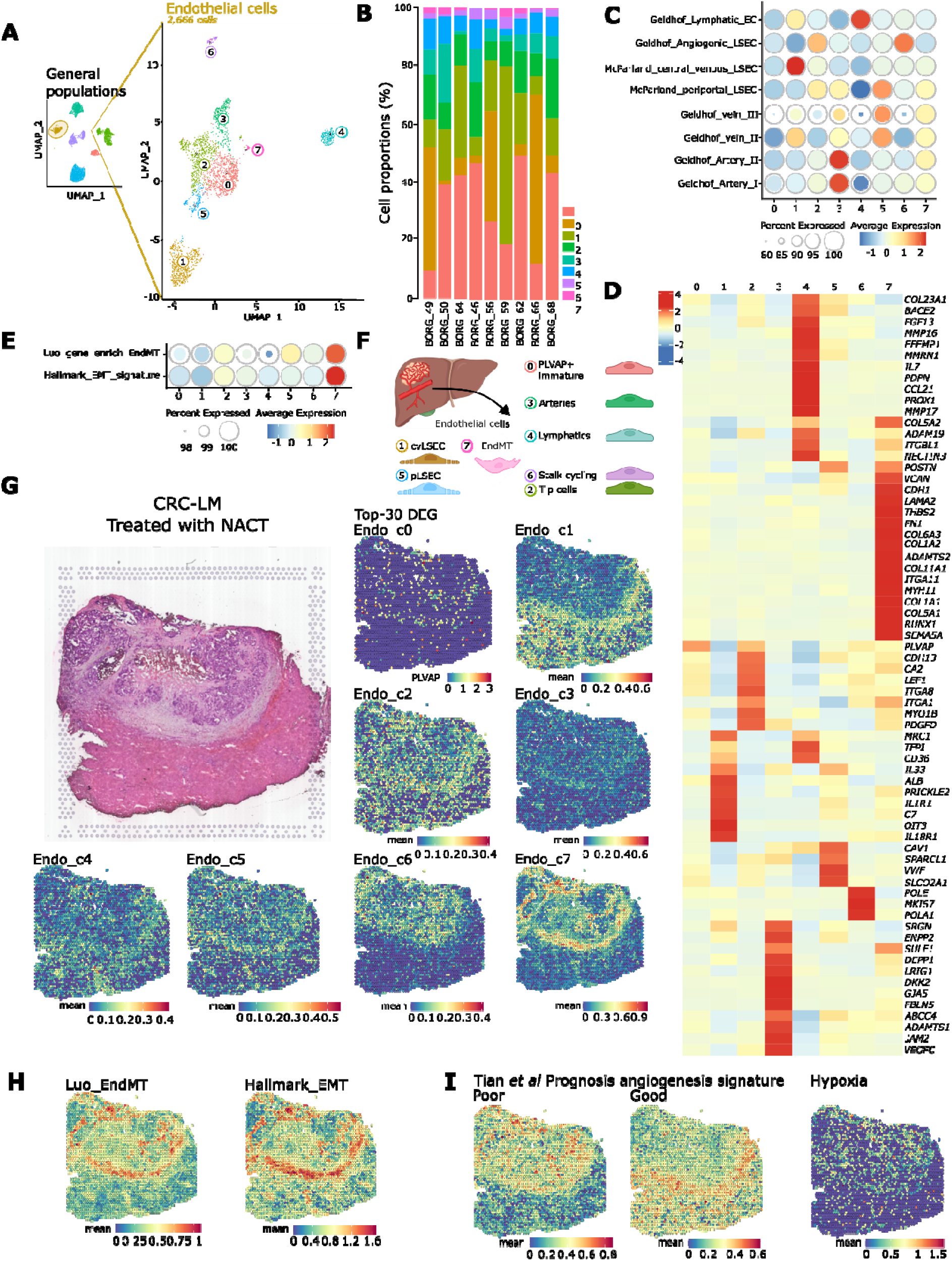
Characterization of the endothelial compartment of LM-CRC. (A) UMAP representing the endothelial clusters obtained by reclustering the endothelial main cellular compartment. (B) Barplot representing the patient’s contributions for each endothelial cluster. (C) Dot plot representing the enrichment of endothelial populations signatures across endothelial clusters. Dot size is for percent of cells expressing the signatures while color scale is for relative expression intensity. (D) Heatmap representing the expression of DEG across endothelial clusters used for their annotation. Color scale is for relative expression intensity. (E) Dot plot representing the enrichment of EndMT and EMT signatures across endothelial clusters. Dot size is for percent of cells expressing the signatures while color scale is for relative expression intensity. (F) Graphical summary of the CRC-LM endothelial cell composition and annotations. (G) Projection of the expression of endothelial clusters metagenes (top-30) on a spatial transcriptomic data from CRC-LM treated by NACT. Color scale is for absolute expression intensity. (H) Projection of the expression of EndMT and EMT signatures on a spatial transcriptomic data from CRC-LM treated by NACT. Color scale is for absolute expression intensity. (I) Projection of the expression of poor and good prognosis angiogenesis signature and hypoxia signature on a spatial transcriptomic data from CRC-LM treated by NACT. Color scale is for absolute expression intensity.

Several populations retained transcriptional features of specialized hepatic or mature vascular endothelial cells. Endo_c1 expressed canonical markers of liver sinusoidal endothelial cells (LSEC), including *OIT3* [45], and was preferentially enriched for a central-vein (cv) LSEC signature. In addition to its sinusoidal identity, this population expressed genes involved in cytokines signaling (as well as Endo_c4), complement activation, and macrophage recruitment. Endo_c5 similarly displayed sinusoidal features but was preferentially associated with periportal (p) LSEC and venous signatures. By contrast, Endo_c3, exhibited canonical arterial endothelial markers and corresponding signature enrichment, whereas Endo_c4 was characterized by lymphatic endothelial-cell markers and lymphatic-specific transcriptional signatures. Endo_c4 also shared cytokine-related programs with Endo_c1. Together, these populations indicate that specialized vascular identities characteristic of the liver remain represented within metastatic tissues (Figure 5C).

In parallel, Endo_c2 and Endo_c6 cluster displayed complementary angiogenic programs. Endo_c2 expressed canonical markers of endothelial tip cells, which guide vascular sprouting and branching [46], whereas Endo_c6 was enriched for genes involved in angiogenesis, consistent with a proliferative state involved in vessel sprout elongation. Both populations, along with Endo_c3 and Endo_c4, exhibited growth factor-related programs, supporting active vascular remodeling across multiple endothelial states. Endo_c0 was distinguished primarily by *PLVAP*, a marker associated with fenestrated and immature tumor endothelial cells [47,48], however, the limited number of cluster-specific DEG precluded a more detailed functional characterization of this population.

A distinct state was represented by the Endo_c7 cluster, which was defined by the expression of multiple mesenchymal- and ECM-associated genes. As pericytes had already been identified within the mural compartment (Fibro_c5), we hypothesized that Endo_c7 might represent an endothelial state with features of endothelial-to-mesenchymal transition (EndMT). Endo_c7 showed enrichment for both the EndMT signature described by Luo’s *et al* EndMT and the Hallmark_EMT gene set (Figure 5E), together with expression of *FN1*, which has been previously shown to drive the EndMT [49] (Supplementary Figure 9-10).

Overall, the endothelial compartment comprised sinusoid fenestrated subtypes (Endo_c1/c5), angiogenic tip and proliferating stalk states (Endo_c2/c6), an immature PLVAP^+^ population (Endo_c0), resident endothelial cells (Endo_c3/c4) and a mesenchymal EndMT-like state (Endo_c7) (Figure 5F). We next asked whether these transcriptionally defined populations occupied unique spatial contexts within chemo-treated CRC-LM.

Spatial mapping of endothelial subtypes uncovered a defined relationship between endothelial states and tissue context. The Endo_c1 was predominantly enriched in non-malignant liver parenchyma, consistent with its hepatic sinusoidal identity. Endo_c2, Endo_c4, and Endo_c5 were detected in both tumor and adjacent liver tissues, whereas the proliferative Endo_c6 population was selectively enriched in tumor regions. By contrast, the EndMT Endo_c7 population localized specifically at the tumor margins. Endo_c3 could not be reliably localized, likely due to its low abundance or weak metagene signal. Similarly, spatial localization of Endo_c0 could not be assessed due to its definition by a single DEG (Figure 5G). Consistent with its transcriptional annotation, spatial enrichment of Endo_c7 overlapped with EndMT-associated signatures (Figure 5H). We further examined whether these endothelial states contained prognostic value associated with angiogenesis signatures and hypoxia regions (*PECAM1, SLC2A1, CA9*). The angiogenesis signature associated with adverse outcome was enriched in tumor core, overlapping with EndMT_Endo_c7 and Stalk_cycling_Endo_c6 subtypes. Conversely, the good-prognosis angiogenesis signature was predominantly detected in adjacent liver parenchyma and overlapped with the central-vein LSEC Endo_c1 population. Hypoxia-associated markers were similarly enriched within core tumor regions, spatially overlapping with Endo_c6 and Endo_c7 (Figure 5I).

Together, these findings reveal substantial vascular heterogeneity within CRC-LM and indicate that endothelial phenotypes are closely linked to local tissue context, with proliferative and mesenchymal-like vascular programs primarily associated with hypoxic tumor regions and the tumor-liver interface. These observations support substantial vascular remodeling within the metastatic ecosystem; however, findings should be interpreted with caution given the relative scarcity of endothelial cells and the resolution limits of the spatial transcriptomics data.

### Liver-resident epithelial populations retain distinct metabolic programs

To characterize non-malignant epithelial compartment of the metastatic liver, we reclustered 9,837 hepatocyte and 1,952 cholangiocyte nuclei into nine distinct subtypes (EpiLiver_c0 to EpiLiver_c8; Figure 6A). Most populations were present across patients at variable frequencies, although EpiLiver_c5 and EpiLiver_c8 contained relatively few cells detected in only two patients and were thus removed from subsequent analyses. Two additional samples (BORG_59 and BORG_64) showed marked predominance of EpiLiver_c3 and were excluded from analyses of relative cell abundance to minimize patient-specific sampling bias (Figure 6B). Expression of canonical lineage markers, such as *CFTR* and *KRT7* for cholangiocytes, and *CP* and *ALB* for hepatocytes resolved the remaining populations into cholangiocyte (EpiLiver_c3 and EpiLiver_c6) and hepatocyte (EpiLiver_c0, c1, c2, c4 and c7) lineages (Figure 6C). Hepatocytes were predominant, counting for 31% (EpiLiver_c0), 24% (EpiLiver_c1), 23% (EpiLiver_c2), EpiLiver_c4 and EpiLiver_c7 being present in lower proportion (6% and 1% respectively). EpiLiver_c3 was the predominant subtype of cholangiocytes, counting 14%, while EpiLiver_c6 represented 1% of cells (Figure 6D). Inferred copy-number profiles were largely diploid and clearly distinct from those of the malignant EpiColon populations, further supporting their non-malignant origin (Figure 6E).

**Figure 6:**
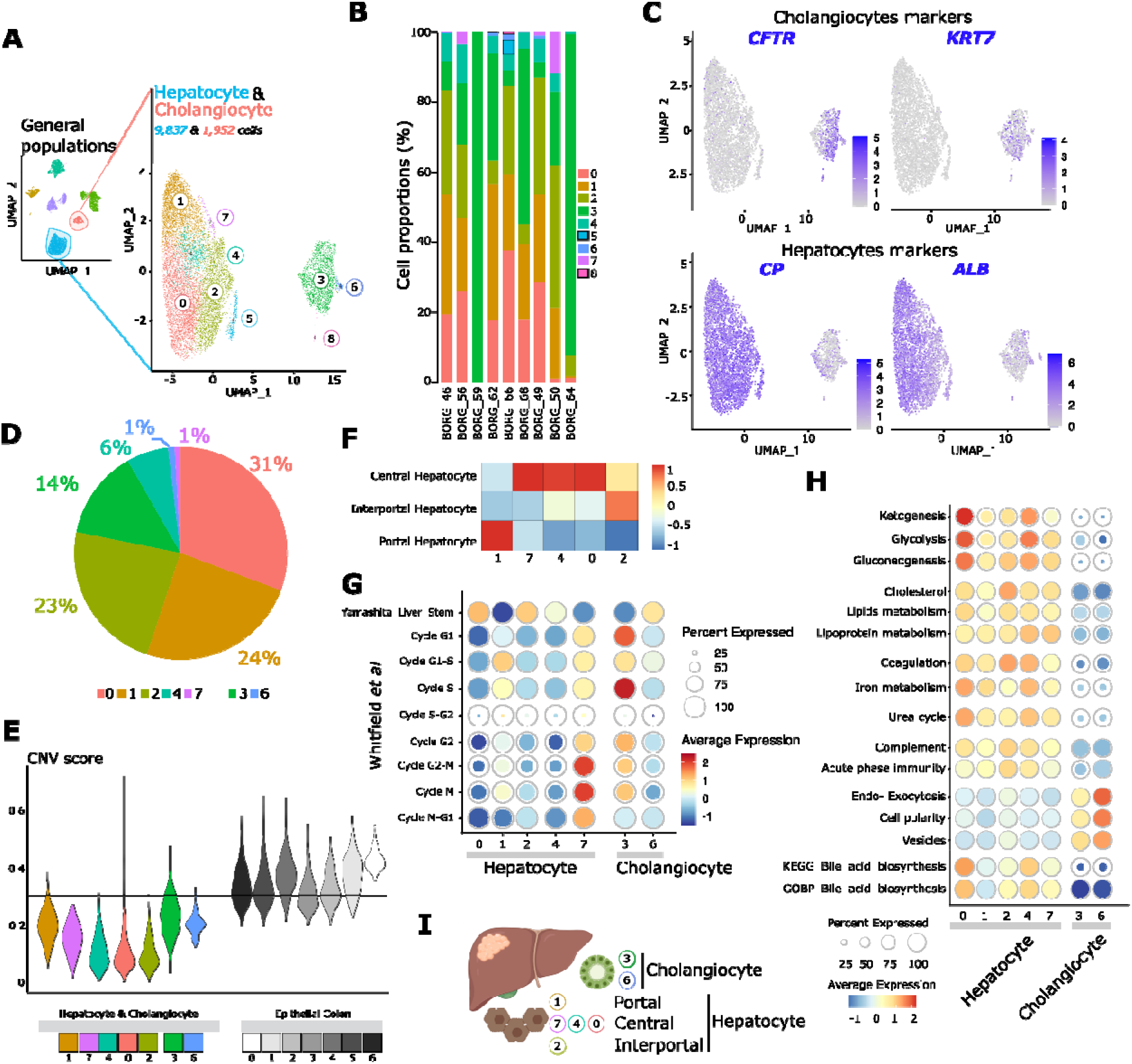
Characterization of the hepatocyte and cholangiocyte compartment of CRC-LM. (A) UMAP of the resident liver clusters obtained by reclustering liver main cellular compartment. (B) Barplot representing the patient’s contribution for each resident liver clusters. Frame highlights the clusters that were removed from further analyses. (C) Dimplot representing the expression of cholangiocytes (CFTR and KRT7) and hepatocytes (CP and ALB) canonical markers across liver resident clusters. Color scale is for absolute expression intensity. (D) Pie chart representing resident liver clusters proportions in all patients except BORG_59 and BORG_64 that were biased toward cholangiocytes enrichment. (E) Violin plot representing the CNV for resident liver clusters compared to epithelial tumor clusters. (F) Heatmap representing the expression of hepatocytes signatures distinguished by their tissue localization, across hepatocytes subtypes. Color scale is for relative expression intensity. (G) Dot plot representing the enrichment of cell cycle signatures and liver stem signature across resident liver clusters. Dot size is for percent of cells expressing the signatures while color scale is for relative expression intensity. (H) Dot plot representing the enrichment of main gene ontologies excepted from liver cells, across resident liver clusters. Dot size is for percent of cells expressing the signatures while color scale is for relative expression intensity (I) Graphical summary of the CRC-LM non-malignant epithelial liver cell composition and annotations.

We next examined whether hepatocyte heterogeneity reflected the physiological zonation of the liver lobule. Application of established hepatocyte zonation signatures identified EpiLiver_c1 as portal hepatocytes, EpiLiver_c0, EpiLiver_c4 and EpiLiver_c7 as central hepatocytes, and EpiLiver_c2 and EpiLiver_c5 as interportal hepatocyte states (Figure 6F). These spatially defined identities were accompanied by differences in proliferative and progenitor-associated programs. EpiLiver_c0, c2, and c4 showed enrichment for stem/progenitor-associated signatures, whereas EpiLiver_c1 and EpiLiver_c7 displayed G1/S- and G2/M-associated cell-cycle programs, respectively. For cholangiocytes, EpiLiver_c3 comprised a mix of cells in S or G1 phase, while EpiLiver_c6 displayed greater enrichment for stem progenitor features (Figure 6G). These findings indicate that the non-malignant epithelial compartment retains established hepatic identities.

Functional analyses of hepatocyte and cholangiocyte subtypes based on DEG and gene ontologies revealed metabolic specialization among the hepatocyte populations (Figure 6H and Supplementary Table 11). Analysis of energy metabolism revealed an enrichment of genes involved in ketogenesis, glycolysis and gluconeogenesis in Central_EpiLiver_c0 and to a lesser extent in Central_EpiLiver_c4. Cholesterol, lipid and lipoprotein metabolism signatures were enriched in similar intensity in all hepatocyte subtypes. Central_EpiLiver_c4 was enriched in both coagulation and iron metabolism genes, while interportal_EpiLiver_c2 was enriched in coagulation genes and Central_EpiLiver_c0 was enriched in iron metabolism genes. Urea cycle signature was enriched in Central_EpiLiver_c0 and c4, while genes involved in complement cascade and acute phase immunity were enriched in similar intensity by all hepatocytes. Cholangiocytes_EpiLiver_c3 and c6 were enriched in endo/exocytosis, cell polarity and vesicles genes testifying from their epithelium organization and their secretory function. Consistently, bile acid signatures were enriched only in hepatocytes, stronger in Central_EpiLiver_c0 and c4 (Figure 6H and Supplementary Table 11).

Collectively, analysis of the liver parenchyma analysis reveals that the liver-resident epithelial compartment recovered from chemo-treated CRC-LM retains substantial features of normal hepatic organization, including hepatocyte zonation, lineage-specific metabolic functions, and cholangiocyte secretory programs. Additionally, differences in metabolic, proliferative, and progenitor-associated programs indicate functional heterogeneity within the non-malignant parenchyma surrounding metastatic lesions (Figure 6I).

### A fibroblast state enriched in early-relapse metastases is associated with tumor dissemination

Given the substantial heterogeneity observed across the cellular compartments of CR-LM, we next asked whether specific features of the metastatic ecosystem were associated with clinical outcome. We compared tumors ecosystem from patients who experienced local relapse within one year after surgery (early relapse; eR, n=3) with those from patients without early relapse (noeR, n=6). Among the transcriptionally defined cell states, ECM-myCAF/Fibro_c3 showed the most pronounced difference between groups, accounting for 34% of fibroblasts in eR samples compared with 5% in noeR samples (Figure 7A-C). Interestingly, Fibro_c3 was also enriched for the CAF-S1 signature, previously associated with early relapse in breast cancer (Figure 7D), supporting its potential role in promoting disease evolution.

**Figure 7:**
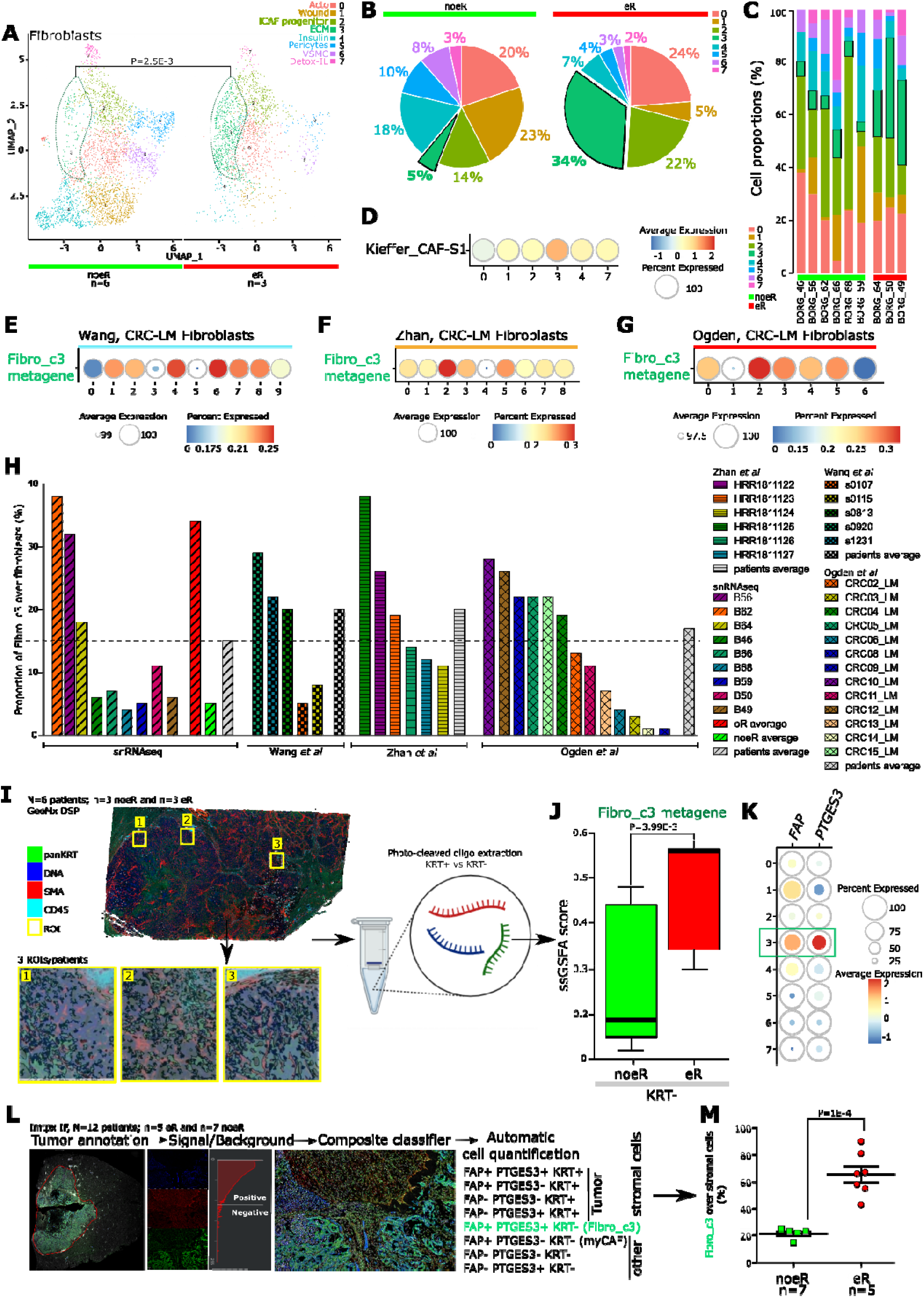
Identification of a fibroblast subtype associated with early metastatic relapse. (A) Segregated UMAP of the fibroblasts clusters between noeR vs eR patients. The green and red color will be attributed all along the figure to these groups respectively. Ovoid frame highlights the significantly enriched fibroblasts cluster. n: effective, P: P-value. (B) Pie chart representing fibroblasts clusters proportions differentially between noeR and eR patients. Frame highlights the proportion of the significantly enriched fibroblasts cluster. (C) Barplot representing the patient’s contribution for each fibroblast cluster. Frame highlights the proportions of the significantly enriched fibroblasts cluster. (D) Dot plot representing the enrichment of CAF-S1 signature across fibroblasts clusters. Dot size is for percent of cells expressing the signature while color scale is for relative expression intensity. (E) Dot plot representing the enrichment of Fibro_c3 metagene across fibroblasts clusters obtained from a CRC-LM scRNAseq (Wang et al). Dot size is for percent of cells expressing the metagene while color scale is for relative expression intensity. (F) Dot plot representing the enrichment of Fibro_c3 metagene across fibroblasts clusters obtained from a CRC-LM scRNAseq (Zhan et al). Dot size is for percent of cells expressing the metagene while color scale is for relative expression intensity. (G) Dot plot representing the enrichment of Fibro_c3 metagene across fibroblasts clusters obtained from a LM-CRC scRNAseq (Ogden et al). Dot size is for percent of cells expressing the metagene while color scale is for relative expression intensity. (H) Barplot representing the proportion of Fibro_c3-like clusters over global fibroblasts population, across datasets (Wang et al, Zhan et al, Ogden et al). Average proportion of Fibro_c3 in eR (red)/noeR (green) groups and global populations (grey) were reported for our snRNAseq dataset. Average proportions of Fibro_c3 in global population (grey) were reported for all the other scRNAseq datasets. Dash line represents the threshold defined by average proportions of snRNAseq dataset. (I) Workflow for GeoMx DSP experiment. (J) Boxplot representing the enrichment of Fibro_c3 metagene in the KRT negative stromal compartment between eR and noeR patients. (K) Dotplot representing the expression of PTGES3 and FAP across fibroblast clusters. Green frame highlights the expression in Fibro_c3 cluster. Dot size is for percent of cells expressing the genes while color scale is for relative expression intensity. (L) Workflow for Fibro_c3 quantification by mtpx IF based on FAP, PTGES3 and KRT markers and whole slide acquisitions. (M) Boxplot representing the proportion of Fibro_c3-like obtained by mtpx IF, over all other stromal cells between noeR and eR groups.

We next evaluated whether a transcriptionally related fibroblast population could be identified in independent single-cell and single-nucleus datasets of CRC-LM. The Fibro_c3 metagene was enriched in three publicly available sc/snRNAseq datasets from Wang_cluster_c4 and Wang_cluster_c6, Zhan_cluster_c2, and Ogden_cluster_c2 (Figure 7E-G), indicating that Fibro_c3-related transcriptional states are reproducibly detected across independent cohorts and profiling technologies. Their relative abundance varied substantially between individual samples in these datasets. For example, using the mean Fibro_c3 abundance in our cohort (15%) as a reference, patients in the Wang (3/5), Zhan (3/6) and Ogden (6/13) datasets segregated into groups with relatively high or low representation of the Fibro_c3 metagene (Figure 7H). Importantly, the absence of annotated clinical relapse information in these datasets precluded testing whether this variation was associated with time to recurrence. Thus, the existence of a conserved Fibro_c3-like state in CRC-LM is supported in external datasets, but its prognostic value should be validated further.

We therefore sought to examine proportions of Fibro_c3 in eR tumors using spatial transcriptomic and protein-based approaches. GeoMx profiling was performed in tumors from 6 patients (n=3 eR and n=3 noeR), with epithelial, immune, and stromal compartments defined using KRT, CD45, and SMA staining and transcriptional profiles obtained from epithelial (KRT^+^) and stromal (KRT^−^) regions (Figure 7I, Supplementary Figure 2). Enrichment analysis of the Fibro_c3 metagene within the stromal compartment revealed a significant increase in eR patients compared with noeR patients (P=3.99 × 10^−3^) (Figure 7J). At the protein level, we selected FAP and PTGES3 among markers preferentially expressed by Fibro_c3 and evaluated their expression by mtpx IF in tumor samples from 12 patients (n=7 eR and n=5 noeR: independent from other patients), together with epithelial tumor cells labeling (KRT^+^). Whole slides images were acquired, tumor regions were delineated by a pathologist, and automatic cell quantification was performed based on these three markers (Figure 7K-L). Consistently, FAP^+^ PTGES3^+^ stromal cells were more abundant in eR patients than in noeR samples (43% to 90% vs 18% to 23%; P=1 × 10^−4^, Figure 7M), supporting the enrichment of a Fibro_c3-like population in tumors associated with early recurrence. Of note, the proportion of Fibro_c3 estimated by mtpx imaging was higher than obtained with transcriptomic analyses, likely reflecting either an intrinsic fragility of this population or suggesting that FAP and PTGES3 may also capture related fibroblast states. Collectively, these analyses support an association between Fibro_c3 abundance and eR, although larger clinically annotated cohorts will be required to establish its independent prognostic value.

We next investigated whether the molecular features and cellular interactions of Fibro_c3 could provide biological insights on this association. Fibro_c3 displayed prominent expression of canonical Wnt/β-Catenin and TGFβ-related pathways, as shown by levels of *WNT7A, WNT2*, and *WNT3A, TGFB2* and *INHBA* which are factors with well-documented roles in epithelial plasticity and EMT [50,51] (Figure 2J-3L). This suggested that Fibro_c3 cells could favor EMT phenotypic plasticity in neighboring cells. Consistent with this possibility, inferred cell-cell interaction analysis between fibroblasts and epithelial tumor cells identified EpiColon_c3 and EpiColon_c1, as candidate interaction partners of Fibro_c3 (Figure 8A). Interestingly, predicted receptor-ligand interactions included the *INHBA-ACVR* converging on the TGFβ signaling pathway (Supplementary Figure 4A-B). These findings are consistent with spatial transcriptomic data showing co-localization of Fibro_c3 with EpiColon_c1 and c3 clusters (Figure 8B) which both displayed features consistent with progressive acquisition of invasive plasticity. EpiColon_c1 showed strong enrichment for EMT-associated signatures (Figure 8C; enrichment in Hallmark metagene, [52]), while the epithelial trajectory analysis described above positioned EpiColon_c3 along an inferred transition toward the invasive/hypoxic EpiColon_c1 state (Figure 2K, Supplementary Figure 3D). To further examine this relationship, we performed pseudotime analysis focused on EMT-related genes between EpiColon_c3 and EpiColon_c1 cells. This analysis revealed a clear activation of EMT-associated programs toward EpiColon_c1 (Figure 8D). Notably, EpiColon_c1 was also enriched for a high-relapse-risk cell signature previously derived from metastatic CRC, further linking this invasive state to programs associated with eR (Figure 8E). Altogether, these data revealed a spatial relationship and signaling crosstalk between Fibro_c3 and malignant epithelial states (eg: EpiColon_c3) displaying enhanced EMT features that may induce epithelial plasticity (eg: EpiColon_c1) and facilitate tumor dissemination.

**Figure 8:**
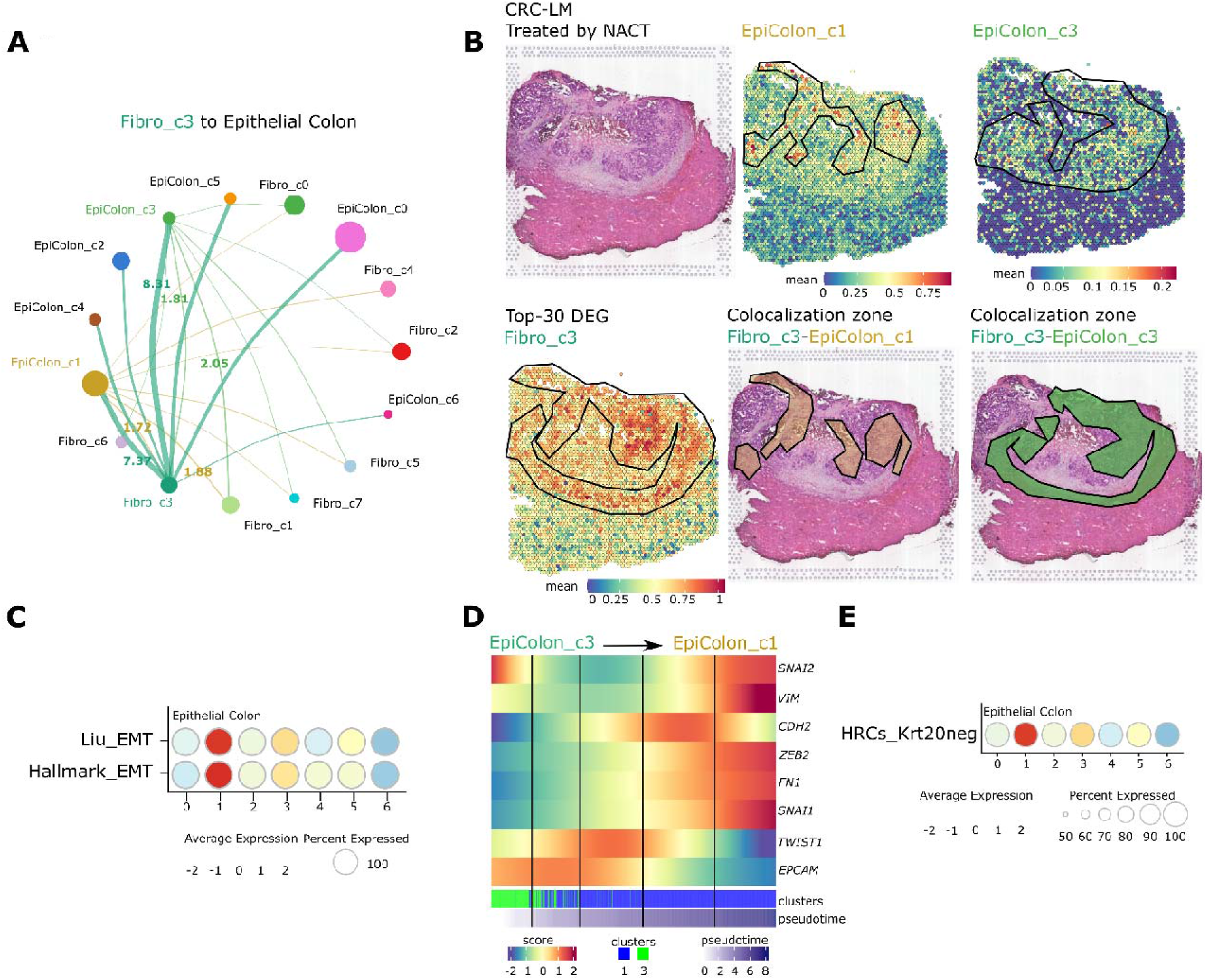
Fibro_c3 & epithelial tumor cells invasion. (A) Network representation of preferential predicted interactions between Fibro_c3 and epithelial tumor cell compartment (and the reverse for two main interactors). Interaction probability is based on the expression of receptor-ligands. Dot represents the relative amount of each cluster. Score are for relative stroke thickness. (B) Projection of the expression of EpiColon_c1, EpiColon_c3 and Fibro_c3 metagene (top-30 DEG) on a spatial transcriptomic data from chemo-treated CRC-LM samples and representation of the colocalization zones between Fibro_c3 and its main interactors. Empty frames represent the region of expression for each cluster of interest, frames with a font color highlights colocalization zone. Color scale is for absolute expression intensity. (C) Dotplot representing the expression of EMT signatures across epithelial tumor clusters. Dot size is for percent of cells expressing the signatures while color scale is for relative expression intensity. (D) Pseudotime gradient map representing the dynamic of expression of EMT-related genes from EpiColon_c3 to EpiColon_c1 clusters. Green to blue scale represents the epithelial differentiation, dark-blue scale represents the pseudotime period. Color scale if for relative expression intensity. (E) Dotplot representing the expression of HRC-KRt20neg signature across epithelial tumor clusters. Dot size is for percent of cells expressing the signature while color scale is for relative expression intensity.

Because dissemination also requires interaction with the vascular compartment, we next examined predicted communication between Fibro_c3 and the endothelial populations defined above. Fibro_c3 primarily interacted with the EndMT-like Endo_c7 subtype (Figure 9A). Notably, Endo_c7 cells from eR samples displayed stronger enrichment for EndMT-associated programs and reduced representation of differentiated vascular features compared with noeR samples (Figure 9B). Predicted Fibro_c3-Endo_c7 communication involved *VEGFA/C-VEGFR, JAG-NOTCH, JAM3*, and other signaling axes implicated in angiogenesis and endothelial plasticity (Supplementary Figure 4C), while NOTCH1 signaling promoted EndMT transcriptional program in endothelial cells [53,54]. Spatial data confirmed the colocalization of Fibro_c3 and Endo_c7 at the tumor margins and in core tumor regions, overlapping with hypoxic areas and enrichment of the invasive/hypoxic EpiColon_c1 state (Figure 9C). Mtpx IF further supported the close spatial association between Fibro_c3 stromal cells and altered vascular structures displaying characteristics of Endo_c7 (Figure 9D-F). Collectively, these analyses identify Fibro_c3 as a fibroblast state co-existing with malignant epithelial and endothelial populations enriched in plasticity-associated programs and associated with early-relapse CRC-LM. The convergence of Fibro_c3 with EMT-like epithelial states and EndMT-like vascular populations suggests a potential stromal environment permissive to tumor-cell invasion and vascular remodeling (Figure 9G).

**Figure 9:**
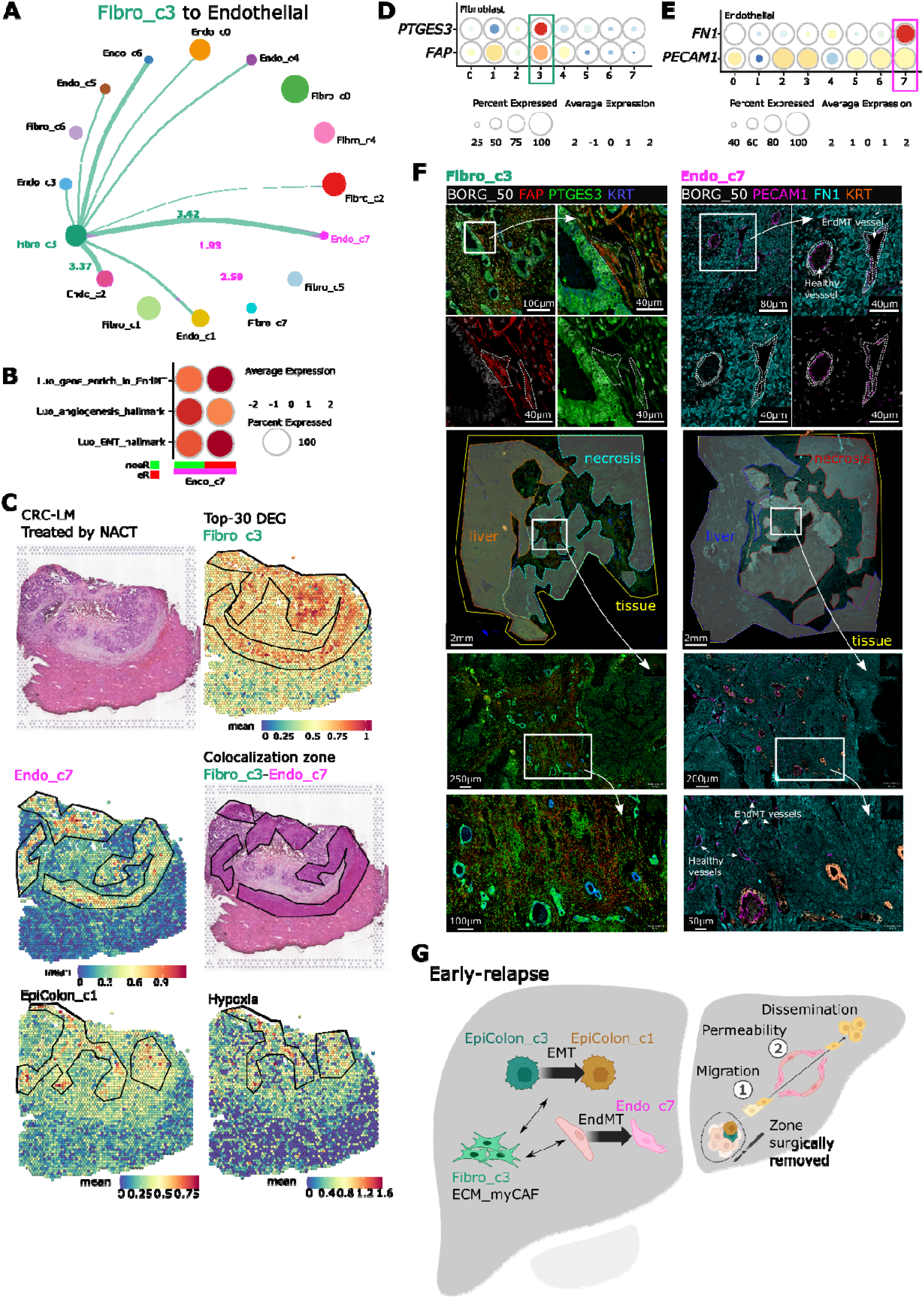
Fibro_c3 & endothelial permeability. (A) Network representation of preferential predicted interactions between Fibro_c3 and endothelial cell compartment (and the reverse for the main interactors). Interaction probability is based on the expression of receptor-ligands. Dot represents the relative amount of each cluster. Score are for relative stroke thickness. (B) Dotplot representing the differential expression of EndMT, angiogenesis and EMT signatures in Endo_c7 cluster from eR (*red*) and noeR (*green*) patients. Dot size is for percent of cells expressing the signatures while color scale is for relative expression intensity. (C) Projection of the expression of Endo_c7 and Fibro_c3 metagene (top-30 DEG) on a spatial transcriptomic data from chemo-treated CRC-LM and representation of the colocalization zone between Fibro_c3 and this main interactor. Empty frames represent the region of expression for each cluster of interest, frames with a font color highlights colocalization zone. Color scale is for absolute expression intensity. (D) Dotplot representing the expression of *PTGES3* and *FAP* across fibroblasts clusters. Frame highlights the enrichment in Fibro_c3. Dot size is for percent of cells expressing the genes while color scale is for relative expression intensity. (E) Dotplot representing the expression of *FN1* and *PECAM1* across endothelial clusters. Frame highlights the enrichment in Endo_c7. Dot size is for percent of cells expressing the genes while color scale is for relative expression intensity. (F) Microscopy images of the colocalization of Fibro_c3 (*FAP and PTGES3* positive cells, *left panels*) and Endo_c7 (*PECAM1 and FN1* positive cells, *right panels*) at the protein level obtained mtpx IF on serial cuts of LM-CRC tissue. Frames and long arrows indicated inserts, small arrows healthy and permeable vessels. (G) Graphical summary of the implication of ECM_myCAF Fibro_c3 in eR by inducing EMT and cell migration in the epithelial tumor cells and EndMT and vessel permeability in the endothelial compartment helping dissemination.

## Discussion

Using an integrated single-nucleus, spatial transcriptomics and proteomics, we characterized the cellular ecosystem of CRC-LM and provided a broad representation of its malignant and non-malignant compartments. Beyond defining the major epithelial, fibroblasts, immune, endothelial and liver-resident cell populations, our analyses revealed substantial functional heterogeneity within each compartment and a distinctive spatial organization of these cellular states in tumor core, tumor margins, and adjacent liver parenchyma.

Most published CRC-LM atlases have relied on scRNAseq, an approach that provides high-resolution characterization of immune populations but can be affected by dissociation-related biases, particularly in fibrotic, necrotic and treatment-exposed tissues. Consistent with this, several studies have focused predominantly on the immune compartment [10,11,36,55–57]. For example, an immunosuppressive switch upon chemotherapy treatment, involving MRC1^+^1CCL18^+^ and SPP1^+^ macrophages, has been described [10]. Other studies have also investigated epithelial and fibroblast heterogeneity [25–28]. Moorman et al characterized the epithelial populations across primary and metastatic CRC and showed that metastatic invasion involves loss of intestinal identity and reprogramming into a highly conserved fetal progenitor-like state before undergoing non-canonical differentiation into divergent squamous and neuroendocrine-like states, a process that is exacerbated in metastasis and by chemotherapy and is associated with poor patient survival [60]. Similarly, Lin *et al* and Zhan *et al* highlighted the involvement of CSC and myCAF in CRC-LM and CRC recurrence, although with limited additional biological characterization [32,58]. Among the most comprehensive studies, Wang et al, profiled primary CRC, CRC-LM and non-malignant tissues and identified multiple epithelial, endothelial, fibroblast and immune populations, providing particularly high granularity within the immune compartment [30]. However, the non-immune clusters were annotated primarily based on their top DEG, without deeper biological characterization or metagene analysis, making direct comparison with our subtypes impossible. More broadly, the authors also demonstrated spatially organized interactions between stromal and immune populations, including MCAM^+^ fibroblasts and CXCL13^+^ cytotoxic T cells through JAG-NOTCH signaling [11]. More recently, by combining both scRNAseq and snRNAseq, Ogden et al identified 14 clusters of epithelial subtypes, 4 of fibroblasts, 4 of endothelial cells, and 30 immune cells. Their fibroblasts compartment included myofibroblasts, pericytes, iCAF and ECM-myCAF whose spatial distribution was assessed by spatial transcriptomic. Consistent with our results, the iCAF cluster was localized both at tumor-liver interface and within the liver, the ECM_myCAF subtype was enriched both at tumor border and within core tumor, while myofibroblasts were spread within tumor core. The latter two clusters also colocalized with myeloid cells subtypes, including C1QC^+^, IL1β^+^ and SPP1^+^ macrophages. A recent study examining hepatic micrometastases from colorectal cancer, using spatial transcriptomics analysis, has highlighted an enrichment in myCAF. The differentially expressed genes in this subtype included *ITGBL1, FAP* and *INHBA*, which are key factors in our Fibro_c3 population [61].

Despite the overrepresentation of immune cells and the relative underestimation of other stromal populations, as well as the limited biological characterization of clusters, these atlases are, to our knowledge, the most comprehensive in literature. Whereas dissociation-based approaches offer particularly deep resolution of immune-cell heterogeneity, our single-nucleus strategy enabled broad recovery of malignant, stromal, vascular and liver-resident epithelial compartments from highly fibrotic and chemotherapy-exposed specimens. We do not interpret that single-nucleus profiling fully reproduces the native cellular composition of the tissue, as each technology introduces distinct technical biases. Our benchmarking suggests that snRNAseq reduces some of the recovery biases associated with tissue dissociation and facilitates characterization of non-immune populations that can be underrepresented in scRNAseq datasets. The integration with spatial transcriptomics and protein-based analyses additionally allowed transcriptionally defined cell states to be placed within their tissue context. Taken together, these approaches should be regarded as complementary, and their combined application will likely provide the most complete representation of CRC-LM ecosystems.

An important feature emerging from our analysis was the organization of malignant epithelial cells along interconnected phenotypic states rather than as completely discrete populations. Stem-like EpiColon_c4 cells were linked through inferred trajectories to early-differentiated states and subsequently to either proliferative populations or an invasive/hypoxic EpiColon_c1 state. The progressive enrichment of EMT-related programs along the EpiColon_c3-to-EpiColon_c1 trajectory, together with the preferential localization of EpiColon_c1 to hypoxic, fibrotic and tumor-budding regions, is consistent with substantial epithelial plasticity within the metastatic microenvironment. These observations are consistent with recent studies describing dynamic and reversible epithelial states during CRC progression and metastasis [62–64]. The stromal compartment showed a similarly complex organization. Rather than aligning with a simple iCAF/myCAF dichotomy, the fibroblast populations identified in our study combined ECM-remodeling, inflammatory, angiogenic, metabolic and immunoregulatory programs. Their spatial organization further suggests functional specialization within distinct stromal niches. The identification of pericytes and VSMC populations as separate mural-cell states also reinforces the heterogeneity of the broader mesenchymal compartment. These stromal populations were closely integrated with immune and vascular programs. In particular, SPP1^+^ tumor-associated macrophages, IL1β^+^ suppressive myeloid cells and immunoregulatory dendritic cells coexisted with lymphoid populations displaying limited cytotoxic activity and exhausted profile, while endothelial cells included both angiogenic populations and an EndMT-like state preferentially located at tumor margins and hypoxic regions. Together, these findings support a model in which immune suppression, stromal remodeling, epithelial plasticity and vascular adaptation are spatially coordinated within the metastatic niche.

After defining the cellular ecosystem, we investigated whether specific cell states were associated with clinical outcome. Despite the limited number of patients, Fibro_c3 showed a marked enrichment in tumors from patients who experienced local relapse within one year after surgery. Importantly, transcriptionally related Fibro_c3-like populations could be identified in three independent sc/snRNAseq datasets, including the studies by Wang *et al*, Zhan *et al* and Ogden *et al*, supporting the existence of a conserved fibroblast state across CRC-LM cohorts and profiling technologies. The relative abundance of this population varied substantially among individual metastases in these datasets. However, the lack of detailed relapse information prevented assessment of whether this heterogeneity was associated with clinical outcome. These external datasets therefore support the biological reproducibility of Fibro_c3; however, its prognostic significance requires validation in independent cohorts.

Beyond this preliminary clinical association, our analyses provide a biological framework that may explain why Fibro_c3 is enriched in tumors with early recurrence. Fibro_c3 expressed WNT/β-catenin- and TGFβ-related mediators and showed strong predicted interactions with EpiColon_c3 and the invasive/hypoxic EpiColon_c1 population. These epithelial states colocalized with Fibro_c3 in spatial transcriptomic data, and pseudotime analysis revealed progressive activation of EMT-related genes along the inferred transition from EpiColon_c3 toward EpiColon_c1. EpiColon_c1 was also enriched for a high-relapse-risk cell signature derived from metastatic CRC [64]. Together, these observations are consistent with a model in which Fibro_c3 contributes to a local signaling environment permissive to epithelial plasticity and tumor-cell dissemination. Interestingly, Fibro_c3 also showed prominent predicted interactions with the EndMT-like Endo_c7 population through VEGF-VEGFR, JAG-NOTCH, JAM3 and related signaling axes. Endo_c7 showed stronger EndMT-associated programs in early-relapse samples and colocalized with Fibro_c3 at tumor margins and within selected tumor-core regions, where both populations overlapped with hypoxia and invasive epithelial states.

Several limitations should be considered when interpreting these findings. First, the discovery cohort was small, comprising nine CRC-LM specimens and only three patients experiencing early relapse. This limits statistical power and precludes robust adjustment for clinical, molecular and treatment-related variables that could influence recurrence. The association between Fibro_c3 and early relapse should therefore be regarded as exploratory. Second, snRNAseq reduces several biases associated with tissue dissociation but introduces its own limitations, including reduced detection of cytoplasmic transcripts and incomplete recovery of particular populations such as granulocytes. In addition, the specimens were obtained after systemic chemo-treatment, making it difficult to distinguish features intrinsic to CRC-LM from states induced by therapy. These hypotheses will therefore require validation in larger clinically annotated cohorts and functional experimental models.

In summary, our study provides an integrated view of the cellular and spatial heterogeneity of chemotherapy-treated CR-LM. The data reveal extensive specialization across malignant and non-malignant compartments and identify spatially organized interactions among epithelial, stromal, immune and vascular populations. Among these, Fibro_c3 was identified as a mesenchymal state enriched in early-relapse tumors and associated with epithelial and endothelial programs, suggesting a link between stromal organization and metastatic dissemination and recurrence.

## Material and methods

### Patients and samples

The B-ORG study (N°ID-RCB: 2019-A00710-57) was approved by the French National Agency for the Safety of Medicine and Health Products, the national ethics committee Sud-Méditerranée and the institutional review board of the Assistance Publique des Hôpitaux de Marseille. This study was conducted in accordance with the Declaration of Helsinki. In accordance with national legislation on the protection of persons participating in biomedical research, each patient enrolled in the study voluntarily signed an informed consent form. This monocentric prospective cohort was promoted and conducted by the Assistance Publique des Hôpitaux de Marseille. The B-ORG cohort allowed the collection of clinical samples at time of surgery of patients diagnosed with liver metastases of colon adenocarcinoma. Depending on the patient’s general condition and the extent of metastatic disease, perioperative chemotherapy was indicated to complete surgical resection. A total of 43 patients admitted to the North Hospital of Marseille were enrolled between January 2019 and December 2022. Follow-up was organized according to current recommendations for good practice.

Collected fresh CRC-LM were both flash frozen for nuclei extraction and formalin-fixed paraffin-embedded (FFPE) for histology. A total of nine samples were selected for snRNAseq based on tissue quality (necrosis) and the time elapsed between surgery and freezing. Analyzed patients included three who experienced disease relapse within one year after surgery and were classified as the early relapse group (eR group), and six who did not (noeR group). Clinical characteristics of the patients and samples are reported in Supplementary Table 12.

### Single nuclei RNA sequencing

#### Nuclei isolation

A batch of two representative samples, including core tumor and tumor margin, were processed at a time. An average of 20mg of tissue was pre-chopped in smaller sections (1mm^3^). Nuclei extraction was performed using Chromium Nuclei Isolation Kit with RNAse inhibitor (PN-1000494; CG00505; 10x Genomics) according to the supplier’s instructions, except an additional wash added to 10X Genomics’s protocol before debris removal. Nuclei were stained with 4’,6-diamidino-2-phenylindol (DAPI, 0.2µg/µL, Thermo Fisher Scientific) to control their quality and estimate the concentration of nuclei suspension, using light and epifluorescent microscope (X40 magnification) regarding to 10X Genomics recommendations. Isolated nuclei were immediately used for downstream processing.

#### Library

The 10X Genomics Chromium Newt GEM Single Cell 3’ Reagents v3.1 kit (CG000315, 10X Genomics) was used with standard conditions and volume to process nuclei suspensions for 3’ transcriptional profiling. The nuclei suspension volumes were calculated for target cell recovery of between 8,000 and 12,000 nuclei for all dissociated samples and loaded on the Chromium according to manufacturer’s guidelines. According to the guidelines of Montpellier GenomiX platform, libraries were pooled and loaded into Miniseq system sequencer (Illumina) to perform a shallow sequencing to visualize the behavior of samples in a sequencing flowcell and to adapt each sample for deep sequencing on Novaseq 6000 Illumina sequencer (Read 1 = 28bp, i7 index=10bp, i5 index=10bp, Read2=90bp).

#### Single-nuclei RNA sequencing data quality control and processing

The resulting intensity files were demultiplexed and then aligned to the human genome, version hg38, using 10X Genomics Cell ranger v3.0.1 software package. scRNAseq analysis was performed using the R package Seurat v4.3.0.1. The nine individual samples were integrated into a single dataset. Cells with more than 10% mitochondrial percentage and 35% of contamination (DecontX) were removed. The ambient RNA profile was removed using SoupeX algorithm [65]. Data were normalized and scaled. We identified the top 2,000 highly variable features using the “vst” selection method. Principal component analysis was then performed, and the first 50 principal components were retained for downstream analysis. This resulted in a final dataset of 31,524 filtered cells.

Cell clustering was performed using the Louvain algorithm with a resolution parameter of 0.5. For visualization, we used UMAP. A two stages clustering strategy was used, in which cells were first annotated into six major cellular populations (immune, endothelial, epithelial tumor cells, fibroblasts, cholangiocytes, and hepatocytes), based on the expression of canonical markers in DEG. Stressed (high mitochondrial percentage) and contaminating cells were removed. Major cellular populations were then partitioned into subtypes deeply characterized looking for the expression of canonical markers, biological pathways, signature enrichment, metabolic profile etc. We distinguished epithelial malignant cells from normal diploid cells based on their copy number profile using InferCNV. Single Cell trajectories and pseudotime were inferred with Monocle 3 and Slingshot. Intercellular communication analysis was done with CellChatDB; the scores mentioned in network plot are for stroke thickness itself representing the probability of interaction.

#### Public datasets

Validation sc/snRNAseq data were obtained from publicly available datasets;

- [dataset] Lin 2024, GSE231559, [32], scRNAseq CRC-LM n=9.
- [dataset] Che, 2021, GSE178318, [66], scRNAseq CRC-LM n=6.
- [dataset] Zhan, 2025, HRA007643, [58], scRNAseq CRC-LM n=12.
- [dataset] Wang, 2023, GSE225857, [11], scRNAseq CRC-LM n=5.
- [dataset] Ogden, 2025, E-MTAB-13651, [12], snRNAseq, CRC-LM n=13.

Each dataset was analyzed independently to bypass batch effects, with the same processing as the authors did.

### Automated multiplexed immunofluorescence

#### Tissue preparation and region of interest selection

Serial 3µm tissue sections were cut using a microtome (HM 340E ThermoFisher Scientific) and loaded on microscope slides for printers (Kilinipath). Tissues were colored by Hematoxylin-Eosin-Safran and scanned using NanoZoomer 2.0-HT (Hamamatsu PhotonicsTM). The scan was visualized using Calopix v5.0.20. One region of interest was selected per tissue, sufficiently large to include the core tumor, the margin and liver front hepatocytes.

#### Staining

Mtpx IF was performed using MACSima™ Imaging Cyclic Staining technology (Miltenyi Biotec), following supplier’s instructions. Tissue and are detailed in Supplementary Figure 1A for each sample. The antibody panel is shown in Supplementary Table 13. Slides were recovered and colored by Hematoxylin-Eosin-Safran and scanned (NanoZoomer 2.0-HT, Hamamatsu Photonics^TM^) for tissue quality control.

#### Images processing

Raw pre-processed tiles were stitched and aligned resulting in the final image using MACS iQ View Analysis Software (Miltenyi Biotec). Nuclei were segmented based on DAPI staining with advanced morphology tool set on minimum diameter 4µm and maximum diameters 20µm, detection sensitivity of 100%, separation force of 60% and smoothing filter Sigma of 1. Cytoplasm was segmented using constraint donut based on multiple channels selected (all membrane signals available), with a detection sensitivity at 100% and donut width at 20µm. The segmentation efficacy, notably for large hepatocytes and dense immune cells, was controlled. The antibody specificity was controlled regarding the expected frequency and morphology. Histograms were optimized for maximal rendering of pictures.

#### Analysis

A unique serial gating strategy was used to determine the cellular composition of CRC-LM analyzed by mtpx IF. Briefly, from all segmented cells, positive CD31 cells, i.e., endothelial cells, were detected. From all the CD31-negative cells, epithelial cells, *i.e*. cytokeratin-positive cells, were identified. Among the epithelial compartment, hepatocytes were distinguished from cholangiocytes and tumor cell using cytokeratin-7 expression, and the hepatocytes commercial marker (Miltenyi Biotec). We also assessed the proportion of proliferating tumor cells (KI67^+^). From all cytokeratin-negative cells, we looked at the expression of fibroblasts markers. We determined the number of cells with the different staining patterns: VIM^+^ ACTA2^+^, VIM^+^ PDPN^+^, VIM^+^ ACTA2^−^PDPN^−^, VIM^−^ACTA2^+^, and VIM^−^PDPN^+^. For each, we controlled and excluded all cells expressing an immune cell marker to guarantee the fibroblasts purity. We do not consider the VIM^−^ACTA2^−^PDPN^−^Immune^−^population because we could not guarantee the fibroblast nature in lack of additional markers, even if the morphological aspect suggested a mesenchymal phenotype. From all cytokeratin-negative cells, we looked at the lymphocyte amount (CD3^+^) and more precisely we assessed the presence of helper T cells (CD4^+^), cytotoxic T cells (CD8^+^), including activated ones (GMZB^+^), regulatory ones (FOXP3^+^), proliferative ones (KI67^+^) and exhausted ones (CD279^+^), and natural killer cells (CD56^+^). From all the CD3-negative cells, we assessed the presence of mast cells (Mast Cell Tryptase^+^) and neutrophils (CD15^+^). For the myeloid compartment, we assess the presence of type 1 macrophages (CD68^+^), type 2 macrophages (CD163^+^) and double positive (Kupffer Cells) but also of dendritic cells (cDC1^+^). Finally, B cells (PAX5^+^) and plasma cells (CD138^+^) were identified. For each marker, either the histogram or the scatter plot was used, and the positivity threshold was determined regarding the signal intensity and morphology. Artefacts were excluded from the analysis.

### Manual multiplexed immunofluorescence

#### Tissue preparation

Serial 3µm tissue sections were cut with a microtome (HM 340E ThermoFisher Scientific) and mounted in an oriented manner onto TOMO® Adhesion Microscope Slides (Matsunami GlassSlides), dried at room temperature before being incubated overnight at 56°C incubator for a first step of dewaxing. Slides underwent a 10-minute-long wash of Histolemon® (CARLO ERBA) to complete dewaxing, then a progressive ethanol bath to rehydrate the tissue. Slides underwent heat-induced epitope retrieval in Tris-EDTA buffer (pH=9.0) following PT Link procedure (Dako Agilent).

#### Staining

To quench FFPE-tissue natural autofluorescence, slides were incubated in a solution of Sudan Black B 0.3% (Sigma Aldrich) ethanol 70% for 15 minutes. Slides were washed with Tween20 0.02% phosphate buffer saline (PBS) 1X for 15 minutes then PBS 1X for 5 minutes. Tissues were permeabilized with Tritonx100 0.5% PBS 1X for 15 minutes and saturated with Normal Horse Serum 5% PBS 1X for 30 minutes. Seven-plex panel manual direct immunofluorescence was performed using the following antibodies (Supplementary Table 14-15) and Sytox (1:10 000 dilution H_2_O, Miltenyi Biotech, Germany). Antibodies mix was incubated 1 hour at room temperature and obscurity. Slides were washed several times by serial baths with saturation solution, then PBS 1X and finally with miliQ water before being slipped with Prolong antifade mountant (ThermoFisher Scientific).

#### Image acquisition

A minimum of 3 ROI per tissue was imaged with confocal microscope (LSM800, Zeiss) at X25 immersion magnification using appropriate exposure times and 2×2 tile scan and then analyzed with automatized software (Zen Black, version 3.3.89.0000). Complementary acquisitions of the complete tissue were performed using a fluorescent scanner (Nanozoomer s60 v2 Digital slide scanner, Hamamatsu) to determine ROI repartitions within the tissue (core tumor or margin) and to guarantee the representativity of acquisitions.

#### Cell quantification

Fibro_c3 scans were processed using QuPath-0.5.1. The core tumor was annotated by an expert pathologist. Positive cells for each labeling (FAP, P23 and CK) were detected using “positive cell detection” tool (default parameters completed by nucleus minimum area set on 2µm and staining intensity set on 15 optical density cellular mean, single stain threshold were adjusted regarding optimal signal to background ratio; optical density cellular mean was set on 25 for P23, on 4 for CK and on 15 for FAP). Single cell classifiers were merged into a composite classifier which was then applied to the whole tissue slide. The percentage of Fibro_c3 (FAP^+^ P23^+^ CK^−^) was determined regarding the stromal cells (i.e., calculated from the total number of cells of the core tumor subtracted from the epithelial CK^+^ cells). Statistical test was a non-parametric T-test.

### Spatial transcriptomics

We performed GeoMx Digital Spatial Profiler (DSP) analysis on samples from 6 patients (respectively n=3 patients in eR (patient B2, B46 and B37) and n=3 in noeR groups (patient B20, B30 and B32), Supplementary Figure 2) according to the GeoMx™ DSP Technology Access Program service of NanoString® (Seattle). Main cell compartments were identified based on fluorescent morphological markers; DAPI (DNA), αSMA (fibroblasts), CD45 (immune cells) and pan-cytokeratin (CK) (epithelial/tumor cells). A pathologist selected the tree most representative and complete ROI (600 by 700 µm) for each patient (total N=18 ROI). Probes were collected from CK positive/negative area of illumination, then sequenced on an Illumina NovaSeq 6000 instrument. Raw target count matrix from GeoMX DSP were pre-processed with default Q3 normalization using the NanoString GeoMX Analysis Suite online tools and imported into R. Processed data were log2-transformed and quantile-normalized to correct for technical biases (limma package). The enrichment of clusters metagene obtained by snRNAseq was evaluated on appropriate cell compartment.

We also analyzed public spatial transcriptomics data using SpatialTME (yelab.site) and ST-liver1 to 4 available sample from Wu *et al* study [67]. Signatures or top 30 differentially expressed genes lists were loaded in gene expression module to assess tissue localization.

## Ethics approval and consent to participate

Our study systematically collected the informed patient’s consent to participate, and the ethics and institutional review board were obtained (see Methods). The study was approved by our institutional review board.

## Availability of data and materials

Single-nuclei RNA sequencing data are available in GSEA portal under accession number.

## Author’s contributions

Conception and design, E.M., D.J.B., F.B.; methodology, E.M., M.P., A.G., P.F.; validation, E.M.; formal analysis, M.P., E.De., C.G., A.G., P.F.; resources, G.L., B.d.R., E.Do; data curation, M.P.; writing - original draft preparation, M.P.; writing - review and editing, E.M., M.P.; supervision, E.M., D.J.B. All authors have read and agreed to the published version of the manuscript.

## Acknowledgment

The authors wish to thank the IPC/CRCM Experimental Pathology Platform (ICEP) and MICS platform for their technical support as well as the “Centre de Ressources Biologique” from the AP-HM.

## Supplementary Figures

**Supplementary Figure 1:**
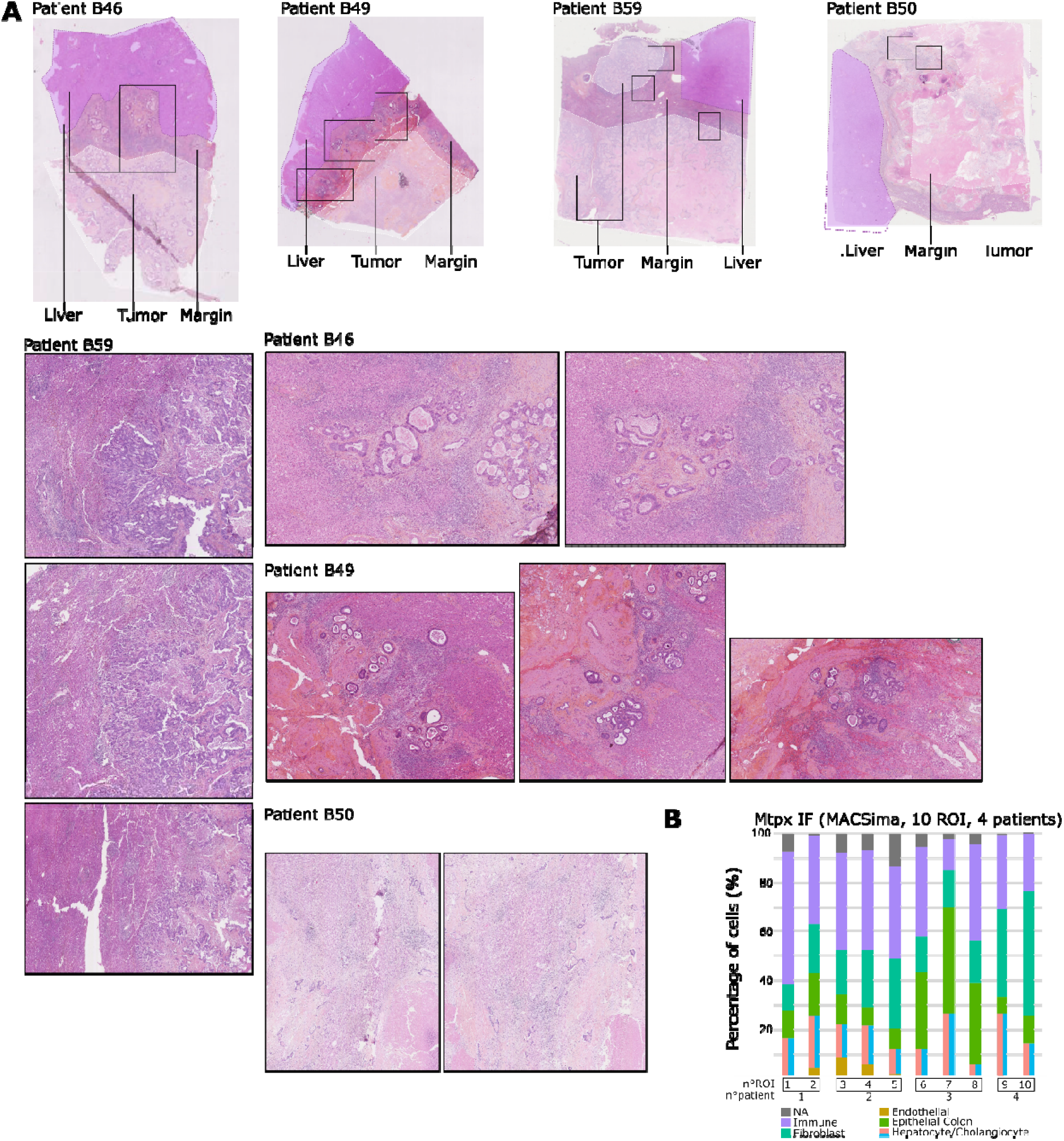
(A) CRC-LM tissue and region of interest analyzed by MACSima automated mtpx IF. Hematoxylin, eosin and safran coloration of the four CRC-LM tissues. Liver, tumor and margin are delineated by dash lines regions; tumor in white and liver in purple. Regions of interest are represented by frames. (B) Barplot representing the percentage of cells for the main cellular compartment obtained by automatic mtpx IF across patients. ROI; region of interest, NA; non-attributed, n; effective.

**Supplementary Figure 2:**
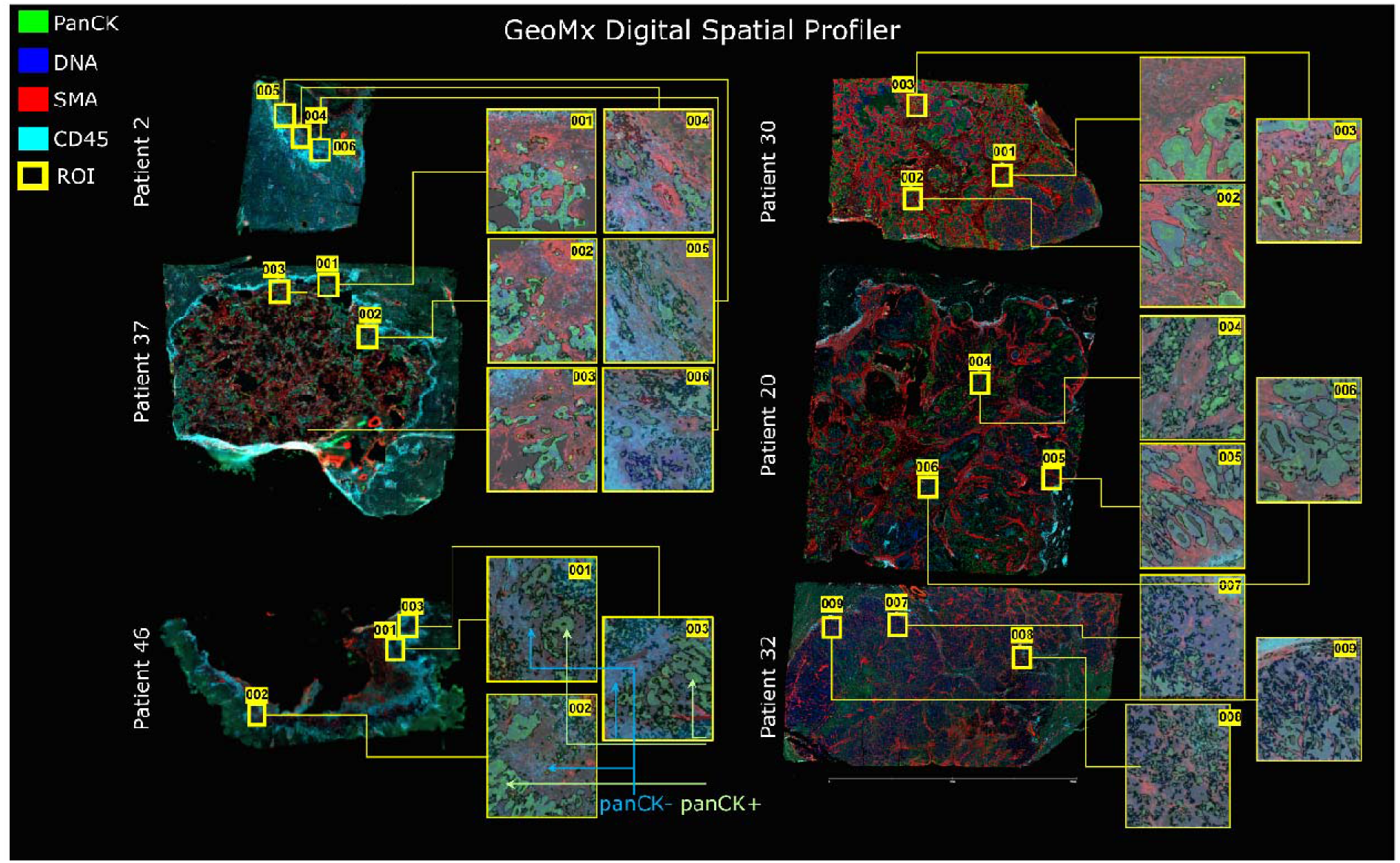
Detail of CRC-LM tissues and ROI analyzed by GeoMx spatial transcriptomic. A total of 6 CRC-LM samples were analyzed; fibroblast, immune and epithelial compartment were labeled (respectively SMA, CD45 and pan-CK) to appreciate cell composition and distinguish transcript from stromal (panCK-) to epithelial (panCK+) compartment. Transcript were recovered by photo-cleaved oligo extraction.

**Supplementary Figure 3:**
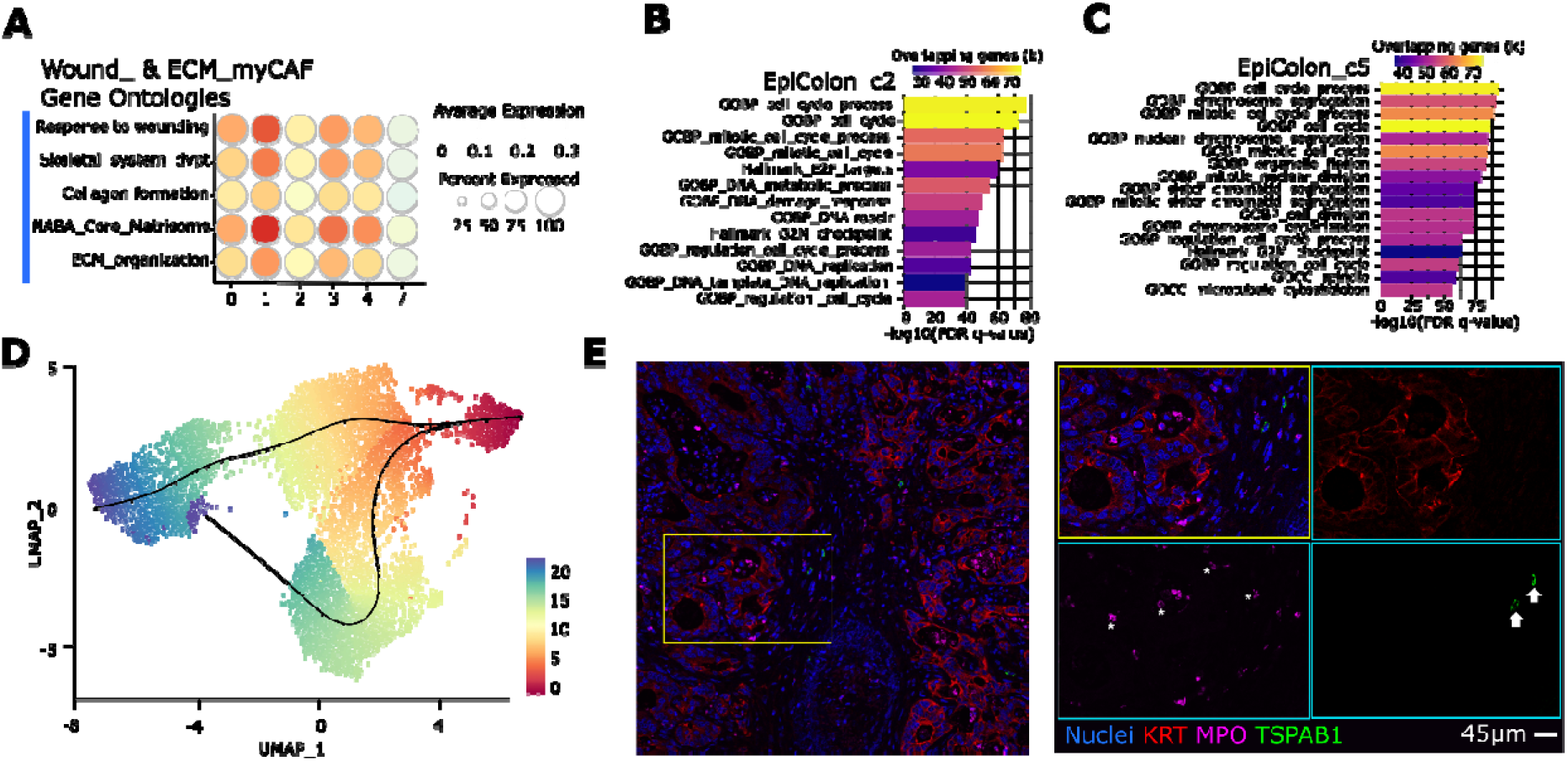
Complement for CRC-LM ecosystem characterization. (A) Dot plot representing the expression of wound- and ECM-myCAF main gene ontologies across fibroblasts subtypes. Dot size is for percent of cells expressing the gene while color scale is for absolute expression intensity. (B) Barplot representing the top enrichment of gene ontologies (GO and Hallmark) in EpiColon_c2 cluster. Color scale is for overlapping genes. (C) Barplot representing the top enrichment of gene ontologies (GO and Hallmark) in EpiColon_c5 cluster. Color scale is for overlapping genes. (D) Pseudotime representation of the trajectory inference from EpiColon_c4 stem-like cluster (defined as origin) toward other epithelial tumoral clusters using Slingshot method. (E) Microscopy images of the granulocytes obtained by manual multiplexed immunofluorescence in CRC-LM sample. Arrows and stars highlight the cells of interest.

**Supplementary Figure 4:**
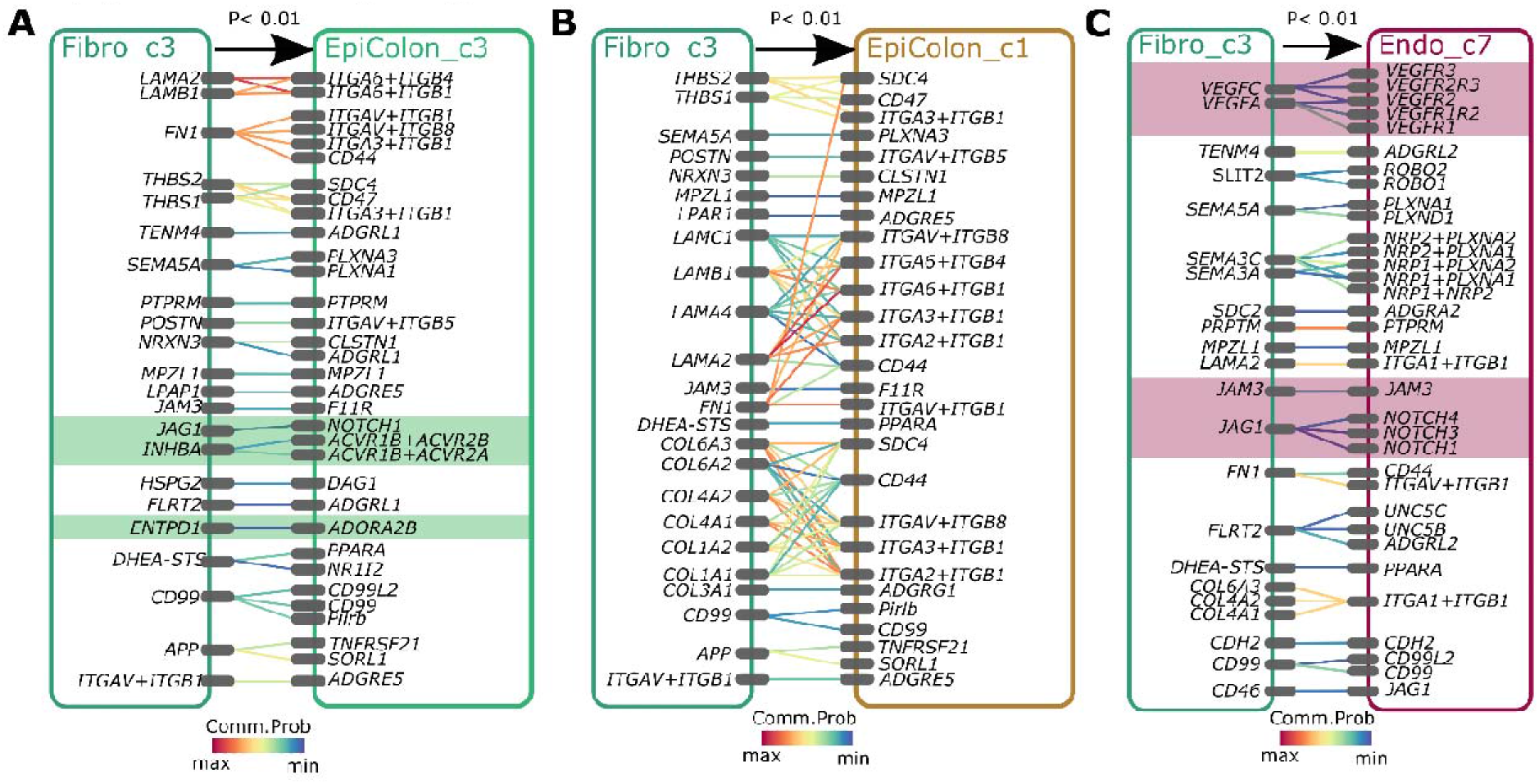
Detail of receptors-ligands involved in the Fibro_c3 main predicted interactions. (A) Detail of receptors-ligands significantly involved in the predicted interaction from Fibro_c3 to EpiColon_c3. The relative probability of interaction (Comm.Prob) is represented by a color scale. B) Detail of receptors-ligands significantly involved in the predicted interaction from Fibro_c3 to EpiColon_c1. The relative probability of interaction (Comm.Prob) is represented by a color scale. (C) Detail of receptors-ligands significantly involved in the predicted interaction from Fibro_c3 to Endo_c7. The relative probability of interaction (Comm.Prob) is represented by a color scale.

